# Metabolic diversity and aero-tolerance in anammox bacteria from geochemically distinct aquifers

**DOI:** 10.1101/2021.09.16.460709

**Authors:** Olivia E. Mosley, Emilie Gios, Louise Weaver, Murray Close, Chris Daughney, Rob van der Raaij, Heather Martindale, Kim M. Handley

## Abstract

**Background:** Anaerobic ammonium oxidation (anammox) is important for converting bioavailable nitrogen into dinitrogen gas, particularly in carbon poor environments. Yet, the diversity and prevalence of anammox bacteria in the terrestrial subsurface – a typically oligotrophic environment – is little understood across different geochemical conditions. To determine the distribution and activity of anammox bacteria across a range of aquifer lithologies and physicochemistries, we analysed 16S rRNA genes, metagenomes and metatranscriptomes, and quantified hydrazine synthase genes and transcripts sampled from 59 groundwater wells distributed over 1 240 km^2^.

**Results:** Data indicate that anammox-associated bacteria (class *Brocadiae*) and the anammox process are prevalent in aquifers (identified in aquifers with sandy-gravel, sand-silt and volcanic lithologies). While *Brocadiae* diversity decreased with increasing DO, *Brocadiae* 16S rRNA genes and hydrazine synthase genes and transcripts (hydrazine synthase, *hzsB*) were detected across a wide range of bulk groundwater dissolved oxygen (DO) concentrations (0 – 10 mg/L). Anammox genes and transcripts (*hzsB*) correlated significantly with those involved in bacterial and archaeal ammonia oxidation (ammonia monooxygenase, *amoA*), which could represent a major source of nitrite for anammox. Differences in anammox community composition were strongly associated with DO and bore depth (and to a lesser extent pH and phosphate), revealing niche differentiation among anammox bacteria in groundwater that was largely driven by water oxygen contents, and not ammonium/nitrite. Eight *Brocadiae* genomes (63-95% estimated completeness) reconstructed from a subset of groundwater sites belong to 2 uncharacterized families and 6 novel species (based on average nucleotide identity). Distinct groups of these genomes dominated the anammox-associated community at dysoxic and oxic sites, further reflecting the influence of DO on *Brocadiae* composition. Six of the genomes (dominating dysoxic or oxic sites) have genes characteristic of anammox (hydrazine synthase and/or dehydrogenase). These genes, in addition to aerotolerance genes, belonging to four *Brocadiae* genomes, were transcriptionally active, although transcript numbers clearly highest in dyoxic groundwater.

**Conclusions:** Our findings indicate anammox bacteria contribute to loss of fixed N across diverse anoxic-to-oxic aquifer conditions, and that this is likely supported by nitrite from aerobic ammonia oxidation. Results provide an insight into the distribution and activity of anammox bacteria across distinct aquifer physicochemisties.

## Introduction

For decades, loss of fixed nitrogen (N) from freshwater environments, including aquifers was exclusively attributed to heterotrophic denitrification [1]. Ammonium (NH_4_^+^) was deemed inert under anoxic conditions [2] and denitrification was considered to be the dominant anoxic sink for inorganic nitrogen [1]. This view was altered when anaerobic ammonium oxidation (anammox), mediated by autotrophic bacteria from a deep-branching class within the phylum *Planctomycetes*, were discovered [3]. Currently, only 10 anammox species have been enriched in culture [4]; none are axenic. Their physiological characteristics and niches are defined using Monod model parameters [5]. Anammox bacteria oxidize ammonium with nitrite (NO_2_^-^) to N_2_ gas without nitrous oxide emissions [6], and are considered chemoautotrophic [7]. Therefore they tend to outcompete heterotrophic denitrifiers in low carbon systems [8]. However, enrichment experiments have demonstrated that some anammox bacteria have the ability to oxidize small organic acids, such as propionate and acetate, with NO_3_^-^/NO_2_^-^ [9], pointing to greater metabolic versatility than previously assumed for these chemolithoautotrophic organisms.

A growing body of research now shows that anammox bacteria play a significant role in the global aquatic N cycle, based on their discovery in diverse aquatic environments, including oceanic oxygen-minimum zones [10,11], meromictic lakes [12], marine subsurface sediments [13], uncontaminated suboxic groundwater [8] and contaminated groundwaters [14–16]. Of these, aquifers harbour the largest freshwater store on Earth, and can be characterized by long groundwater residence times [17], suitable for slow-growing anammox bacteria (with doubling time of 7–22 days) [18]. Moreover, while groundwater often contains low levels of naturally occurring N species, such as nitrate (<0.25 mg/L) [19] and ammonia (<0.2 mg/L) [20], and frequent inputs of N from agricultural sources [21], some aquifers contain little organic carbon [22], potentially limiting heterotrophic denitrification [1]. Aquifers, as oligotrophic environments [22], may therefore represent ideal ecosystems for chemolithoautotrophic anammox bacteria.

Anammox relies on the presence of both oxidized and reduced inorganic N compounds, and subsequently, is highly active in redox transition zones, where N-species fluctuate between oxidation and reduction reactions [23]. Anammox bacteria are therefore found naturally occurring at the interface of aerobic-anaerobic conditions where NO_2_ ^−^ containing water and anaerobic water with NH_4_^+^ mix [24]. N-fluctuation frequently occurs in groundwater due to interactions with surface water, or oxygen penetration from above in unconfined shallow aquifers [25]. Elevated groundwater tables, which promote oxygenation through interactions with the overlying soil and groundwater recharge, stimulate a variety of anammox bacteria[16]. Surface water contaminated with nitrogen-species can be a source of NH_4_^+^ in aquifers. This ammonium can be partially absorbed by clay minerals [26] or oxidised to NO_2_^-^ or NO_3_^-^ by nitrification [1]. NO_2_^-^ produced by aerobic ammonia-oxidizers, can serve as an electron acceptor in anammox [27]. While anammox bacteria have been observed previously in groundwaters [8], their prevalence and genetic diversity is unclear. Nor is it known how they are impacted by the heterogeneous chemical conditions found in aquifers.

To provide insights into anammox ecology and activity across a diverse range of groundwater ecosystems, we collected groundwater from 59 sites over 4 geographic regions across New Zealand. To the best of our knowledge, this is the largest survey of anammox communities in groundwater to date, encompassing a range of characteristics including variations in nitrogen-species, organic carbon and dissolved oxygen concentrations. By reconstruction and metabolic characterisation of diverse novel *Brocadiae* genomes, and by quantifying the distribution, abundance and activity of anammox bacteria across distinct groundwater conditions, we reveal the diverse ecological niches of these bacteria in groundwater. Results provide evidence in support of anammox as an important biological process in aquifers and a major N-sink [16].

## Materials and methods

### Study sites and sample collection

Groundwater was collected from 59 wells spanning 10 aquifers in Waikato, Wellington and Canterbury regions of New Zealand (Supplementary Table S1), from several aquifer lithologies: alluvial sandy gravel (×71 samples), sand-silt (×1), basalt (×1), shellbed (×1), peat (×1) and ignimbrite (×6). Wells were purged (∼3–5 bore volumes), then 3–90 L of groundwater or 0.5–15 L of biomass-enriched groundwater were filtered on-site. Biomass-enriched groundwater was collected from 8 sites in Canterbury (Table S1) directly following standard groundwater collection, and low-frequency sonication (2.43 Kw) for 2 minutes to detach biofilms and aquifer particles [28]. Biomass was captured onto mixed cellulose ester membrane filters (1.2 µm pore size pre-filter over a 0.22 µm filter), using a 142 mm stainless steel filter holder (Merck Millipore Ltd, Cork, Ireland). Both filters were immediately submerged in RNA Later (ThermoFisher Scientific, Waltham, MA, USA), transported on dry ice and stored at -80 °C.

### Chemical analysis of water samples

Dissolved oxygen (DO), water temperature, pH, oxidation-reduction potential (ORP) and specific conductance (SPC or conductivity) were measured on-site using a flow-through cell and field probes (YSI EXO sonde 2, YSI PRO + and YSI ProDSS, Yellow Springs, Ohio, USA). Samples were categorized by DO concentration as follows: anoxic (0 mg/L), suboxic (<0.3 mg/L), dysoxic (0.3–3 mg/L) and oxic (>3 mg/L) [29]. Unfiltered groundwater was analysed for P, N, C, S, Mn and alkalinity at Hill Laboratories (Hamilton, New Zealand) (Supplementary Information). To assess denitrification/anammox, excess N_2_ gas was quantified using methods by Martindale et al. [30]. Briefly, dissolved Ar, Ne and N_2_ were measured by a standard curve via a system comprising two detectors, a pulsed discharge helium ionization detector (Valco Instruments D-4-I-SH14-R), a thermal conductivity detector (Shimadzu TCD-2014) and ultra-high purity helium (He) gas. Excess N_2_, attributable to denitrification/anammox, was determined by comparison to inert Ar and Ne gas concentrations.

### Nucleic acid extraction and sequencing

#### Nucleic acid extraction

RNA and DNA were extracted using RNeasy PowerSoil Total RNA and DNeasy PowerSoil Pro kits (Qiagen, Valencia, CA, USA), respectively, with nuclease-free glycogen added to aid RNA precipitation (0.1 µg/µL final concentration, Roche diagnostics, Basel, Switzerland). DNA extractions used 0.14–0.89 g of crushed filter (1–47 extractions per sample). For RNA, 2.12–3.90 g was used per extraction (1 extraction per sample). RNA was DNase treated using the TURBO DNA-*free*™ Kit (“rigorous” protocol; Invitrogen, Carlsbad, CA, USA). DNA removal was verified via 16S rRNA gene amplification (as below, but over 55 cycles) and gel electrophoresis. Replicate DNA extractions were pooled and concentrated using sodium acetate (0.3M final concentration) and ethanol (2x volume) precipitation with 0.1 µg/µL glycogen (Roche diagnostics) via ethanol precipitation. RNA extractions were concentrated using the Zymo RNA Clean and Concentrator-5 Kit (Zymo research, Irvine, CA, USA).

High molecular weight DNA for whole genome shotgun (WGS) sequencing was verified via 1% agarose gel electrophoresis. Nucleic acids were quantified with a Qubit 3.0 fluorometer (ThermoFisher Scientific, Waltham, MA, USA) using dsDNA HS and RNA HS assay kits, and quality checked using a NanoPhotometer (Implen, Munich, Germany). RNA was checked using an Agilent BioAnalyzer with RNA 6000 Nano and Pico chips (Integrated Sciences, NSW, Australia). Samples with a RNA Integrity Number ≥6 or DV200 >30% (percentage of RNA fragments above 200 nucleotides) were used to quantify transcripts and for transcriptomics. Between 24 pg and 1.1 µg of total RNA was converted to cDNA using Superscript III Supermix (Invitrogen, Carlsbad, CA, USA).

#### Metagenome, metatranscriptome and amplicon sequencing

DNA libraries for 15 samples (gwj01–gwj16) were prepared using the TruSeq Nano DNA Kit (Illumina, San Diego, CA, USA), with a targeted insert size of 550 bp, at the Otago Genomics Facility (University of Otago, NZ), except low-yield sample gwj02, which was prepared with the ThruPLEX DNA-seq Kit (Takara Bio USA, Inc., Mountain View, CA, USA). 2 × 250 bp sequencing was performed using the Illumina HiSeq 2500 V4 platform. RNA libraries were prepared using the Ovation SoLo RNA-Seq System (NuGEN, Redwood City, CA, USA) using custom probes for rRNA depletion at the Otago Genomics Facility, NZ. Custom rRNA probes were designed by the manufacturer using small and large ribosomal subunit sequences reconstructed from the 16 metagenomes with EMIRGE [31] over 40 iterations with clustering at 97% identity and using the SILVA 132 database [32]. Ribosomal sequences generated were used as target sequences in the design of custom AnyDeplete probes using NuGEN’s proprietary algorithm. rRNA and genomic DNA were removed as a part of the library preparation kit, using the custom probes and DNAse treatment respectively. Paired-end 2×125 bp reads were generated from RNA libraries using the Illumina HiSeq 2500 V4 platform.

PCR amplification of 16S rRNA genes used modified Earth Microbiome Project primers EMP-16S-515’F and EMP-16S-806’R primers [33,34] with Illumina Nextera adapters, and MyTaq HS Red Mix (Bioline, London, UK). PCR conditions are described in Table S2. Amplicons were purified using Agencourt AMPure XP beads (Beckman Coulter, Brea, CA, USA). Barcoded libraries, prepared according to Illumina’s 16S Metagenomic Sequencing Library Preparation manual, were loaded with 10% PhiX, for 2 × 250 bp sequencing via Illumina MiSeq with V2 chemistry (Auckland Genomics, University of Auckland, NZ).

### Amplicon processing

Sequences were quality checked using FastQC v0.11.7 [35], and merged using USEARCH v9.0.2132[36]. Sequences were quality filtered using sickle (minimum Phred score ≥30; length ≥200 bp) with another 10 bp of lower quality sequence removed from each end using USEARCH -fastx_truncate [36]. Sequences were dereplicated and clustered at 97% similarity with chimera removal to generate operational taxonomic units (OTUs) using the UCLUST pipeline [37]. OTUs were classified using USEARCH -sintax with the SILVA SSU Ref NR99 database v132 [32]. Non-prokaryotic and singleton sequences were removed before rarefying to 13,393 using QIIME2 v2018.2 [38].

### Quantitative PCR of hydrazine synthase and ammonia monooxygenase genes

Droplet Digital PCR (ddPCR) via the Bio-Rad QX200 platform used 20 µl reactions with 10 µl 2x QX200 ddPCR EvaGreen Supermix (Biorad, Hercules, CA, US), 0.4 µM of forward and reverse primer, 1 µl of DNA or cDNA and nuclease-free water. Primers and PCR conditions for hydrazine synthase (hzsB_396F/hzsB_742R), archaeal ammonia monooxygenase (Arch-amoAF/Arch-amoAR) and bacterial *amoA* (amoA1F/amoA-modR) genes are described in Table S2. Negative controls used DPEC-treated water. Positive controls used gBlocks dsDNA fragments of *hzsB* and *amoA* genes (Integrated DNA Technologies, Coralville, Iowa, US). Data was analyzed using the QuantaSoft software package (Bio-Rad). Threshold values for positive droplets were set based on the amplitude of negative and positive droplets using the positive control as reference.

### Metagenome assembly and genome binning

Adapters were removed from metagenomic reads using Cutadapt [39]. Reads were trimmed with sickle (Phred score ≥30; length ≥80 bp) and quality checked using FastQC v0.11.7 [35]. All samples were individually assembled using SPAdes v3.11.1[40] (--meta, -k 43,55,77,99,121). Samples from the same well (groundwater ± biomass-enrichment) were also co-assembled using identical parameters. Scaffolds ≥2 kb long were binned with MetaBAT2 v2.12 [41], Maxbin v2.2.6 [42] and CONCOCT v1.0.0 [43]. Best-scoring bins per assembly were selected with DAS_Tool v1.1.1[44]. Bins were de-replicated across assemblies using dRep v2.0.5 (average nucleotide identity, ANI > 99%; completeness >50%) [45], and manually refined using *t-SNE* transformation of tetranucleotide frequencies and coverage (https://github.com/dwwaite/bin_detangling). Genome completeness was estimated using CheckM [46]. For genome coverage, trimmed reads were mapped onto de-replicated genomes using bowtie2 v2.3.2 [47] (-n 1 -l 222 --minins 200 --maxins 800 –best). Sample-specific genome relative abundance was calculated by normalizing to library size and highest read count [48]. Estimated genome size was calculated as (bin size – (bin size * contamination))/(completeness) [49].

### Metabolic predictions

Protein-coding gene sequences were predicted using Prodigal v2.6.3 [50] and annotated using USEARCH v9.02132 [36] with -usearch_global (–id 0.5 –evalue 0.001 –maxhits 10), and UniRef100, UniProt [51] and KEGG databases [52]. Hidden Markov Model (HMM) searches were carried out using HMMER v3.3[53] against PFAM [54], TIGRfam [55] databases and databases from Anantharaman *et al*. [56] (HMM individual cutoffs, Table S4). BLASTP (NCBI, National Center for Biotechnology Information) was used to identify predicted protein-coding sequences for Rieske/cytochrome-b systems and the electron transfer module from *Candidatus* Kuenenia stuttgartiensis (similarity >30%, query coverage >70%) (Tables S4-5).

### Genome classification and reconstruction/recovery of 16S rRNA gene sequences

Metagenome-assembled genomes (MAGs) were taxonomically classified using the Genome Taxonomy Database taxonomic classification tool, GTDB-Tk v0.2.1 [57]. 16S rRNA gene sequences were reconstructed from metagenomic data using EMIRGE [31] or SPAdes [40] with identification by Metaxa2 [58] and PATRIC [59]. EMIRGE was used to reconstruct 16S rRNA gene sequences from metagenomes with 97% cluster identity, 80 iterations and the SILVA SSU Ref NR99 database v132 [32].

### Phylogenetic and protein sequence trees

Bacterial core gene alignments (75-114 genes per genome, Table S3) generated from GTDB-Tk were used to construct a maximum-likelihood phylogenomic tree in IQ-TREE (v1.6.9) [60] using ModelFinder [61] best-fit model LG+F+R5, and annotated with iTOL [62]. EMIRGE, Metaxa2 and amplicon sequences were aligned using MUSCLE [63] with default parameters and trimmed to 295 bp using Geneious 11.1.2 (https://www.geneious.com). A maximum-likelihood tree was constructed as with IQ-TREE using ModelFinder best-fit model TIM3+F+I+G4. Protein sequences of hydrazine synthase subunits A, B and C were aligned with MUSCLE and trimmed to remove columns with > 50% gaps using trimAl [64]. A maximum-likelihood tree was constructed as with IQ-TREE using ModelFinder best-fit model WAG+G4.

### Metatranscriptome processing

Adapters were removed from transcriptomic reads, and quality trimmed and checked as described above for metagenomic reads. Residual ribosomal RNA sequences were removed using SortMeRNA v2.1 [65]. Additional checks were carried out to ensure filtered reads were still paired and read pairs were ordered using a repair script from BBMap v38.81 [66]. Filtered transcriptomic reads were mapped to contigs from the set of dereplicated MAGs using Bowtie2 [47] (v2.3.5, --end-to-end --very_sensitive). Read counts were determined using featureCounts [67] (v1.6.3, -F SAF). Singleton reads per gene were removed, and the remaining read counts were normalized to transcripts per kilobase per million reads mapped (TPM) using the following equation: (number of reads mapped to gene)*(1000/gene length)*(1000000/library size) [68].

### Statistical analyses

Analyses were carried out in RStudio (v4.0.3) [69] with the packages: vegan v2.5.6 [70] (for distance-based redundancy analysis, dbRDA) and phyloseq v1.34.0 [71] (alpha diversity analysis). All correlations were Spearman’s rank correlations (*p*-value <0.05 was considered significant).

## Results and discussion

### Prevalence and activity of anammox in chemically diverse groundwaters

#### Anammox bacteria are widespread in groundwater and phylogenetically diverse

All known anammox bacteria exclusively associate with order *Candidatus* Brocadiales [72] in the class *Candidatus* Brocadiae (*Brocadiae*). Based on 16S rRNA gene amplicon analysis of 80 groundwater samples collected across 10 aquifers, *Brocadiae* was present in 60 samples (from 47 wells) across all four regions assessed in this study. Results demonstrate *Brocadiae* prevalence within aquifers comprising unconsolidated materials (sandy-gravel=55/71, sand/silt=1/1), which are a common aquifer type globally [73], and consolidated volcanic rock (basalt/ignimbrite=4/7) (Fig. 1a, Table S1). Furthermore, *Brocadiae* was presence in groundwaters encompassing a range of nitrate-N (0–22 g/m^3^), DO (0–10.6 mg/L), phosphate (0–16.4 g/m^3^) and dissolved organic carbon (DOC, 0–26 mg/L) concentrations (based on bulk groundwater measurements, Table S1). The wide range of groundwater DO concentrations they were detected across is consistent with findings from two pristine carbonate-rock aquifers in Germany [8]. There, anammox was presumed to outcompete denitrification due to low organic carbon, and contributed to 83% of total N loss. Overall, *Brocadiae* represented 0.37% of groundwater prokaryotic communities (3 940/1 071 440 sequences; ∼49.3 sequences ± 202 SD per sample), and ranked 36^th^ in abundance out of all 205 classes of bacteria and archaea. Research has shown that anammox is globally common in soils overlying aquifers, periodically saturated by high groundwater tables [16]. Results here demonstrate anammox-related bacteria are also common in the sediments and rock constituting aquifer saturated zones.

**Figure 1.**
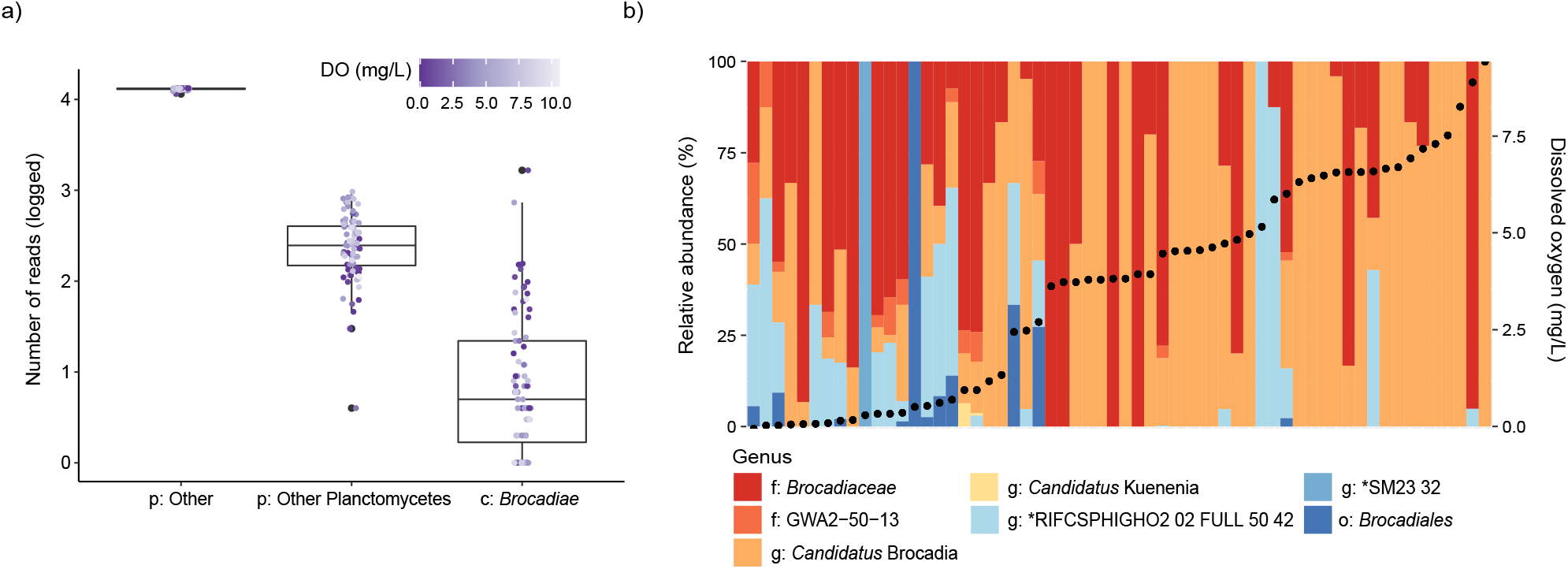
Boxplot and stacked bar graph show the prevalence, relative abundance and diversity of anammox bacteria (*Brocadiae*), based on 16S rRNA gene amplicon data. **a)** Number of OTUs identified as *Brocadiae* relative to other bacteria (p:Other) and non-anammox *Planctomycetes* (p:Other *Planctomycetes*) across 80 samples. Horizontal line represents the median number of OTUs present. **b)** Structure of the anammox bacterial community observed in 60 samples where *Brocadiae* were identified, ordered by DO concentration. Black points = DO content. Bars represent different genera within the class *Brocadiae*. Acronyms: g = genus; f = family; o = order; c = class; p = phylum.

The class *Brocadiae* comprised 37 OTUs (Table S7), of which 42% were assigned to four uncultivated or enrichment-cultivated genera (Fig. S1): *Candidatus* Brocadia (6 OTUs), *Candidatus* Kuenenia (1 OTU), *Planctomycetes* bacterium RIFCSPHIGHO2_02_FULL_50_42 (6 OTUs), and *Planctomycetes* bacterium SM23_32 (3 OTUs). Over half (56.8%) could only be classified at the family (*Brocadiaceae*, 10 OTUs; GWA2-50-13, 9 OTUs) and order (*Brocadiales*, 2 OTUs) levels, highlighting a large degree of novelty among groundwater anammox candidates. Three of these taxonomic groups were strikingly abundant and prevalent, including the 7 most abundant OTUs overall, and representing >90% of all anammox sequences, *Ca*. Brocadia, RIFCSPHIGHO2_02_FULL_50_42, and those classified only as *Brocadiaceae* (Fig. 1b). Together with previous studies, results show that the genus *Ca*. Brocadia is common in terrestrial subsurface environments [8], dominating groundwater anammox communities by up to 80% [74,75]. They were present in 83% of our samples, and comprised 25% of 16S rRNA gene sequences from anammox candidates. The wide distribution of *Ca*. Brocadia across varied groundwater conditions, as determined here, may be attributed to its growth properties (inferred from Monod models) [5]. *Ca*. Brocadia species exhibited the highest maximum specific growth rate among anammox bacteria, and outcompete other freshwater anammox bacteria [5].

#### Brocadiae relative abundance and community composition varies strongly with dissolved oxygen

The overall proportional abundance of *Brocadiae* 16S rRNA gene sequences was significantly and negatively correlated with DO concentrations (*r*=-0.28) and ORP (*r*=-0.27; Table S8), a trend also demonstrated in batch culture experiments [76] and observed in carbonate-rock aquifers [8]. Significant positive correlations were instead found with DOC (*r*=0.29), TDS (*r*=0.23), conductivity (*r*=0.27), DRP (*r*=0.31) and NO_2_^-^ (*r*=0.31, although only 14% of samples had concentrations above the detection limit for NO_2_^-^). TDS, conductivity, DRP and NO_2_^-^ concentrations negativity correlated with DO overall (*p* <0.05). Correlations between *Brocadiae* and other substrates, including those critical for ammonia oxidation (NH_4_^+^ and NO_3_ ^-^), were not significant, indicating that parameters such as DO, ORP, nitrite and DOC are more important factors for controlling the presence of anammox bacteria. Of these parameters, multivariate regression tree analysis, showed that redox potential (ORP threshold 175.9 mV) was the main factor discriminating *Brocadiae* communities (Fig. S2).

Analysis of anammox community diversity by oxygenation regime (anoxic, suboxic, dysoxic and oxic) revealed OTU richness and alpha-diversity (Inverse-Simpson) were significantly higher at the suboxic and dysoxic sites compared with oxic sites (Fig. 2a), suggesting groundwater with low oxygen supports more anammox species. The spatial distribution of different *Brocadiae* taxa was significantly associated with changes in DO and bore depth (db-RDA, Fig. 2b), along with aquifer location and lithology, phosphate, and pH, which collectively explained 16% (R^2^ adjusted) of the variation (Permutation-test, *p* <0.05, permutations=999, Table 8). When considering only DO, dramatic differences in composition were evident (Fig. 1b). Notably, the *Ca*. Brocadia fraction was positively correlated with DO concentrations (*r*=0.46) (Fig. 1b). Three *Ca*. Brocadia and *Brocadiaceae* OTUs comprised up to 30.4—89% of the anammox-associated community in groundwater with >4 mg/L DO (Figs S3-4).

**Figure 2.**
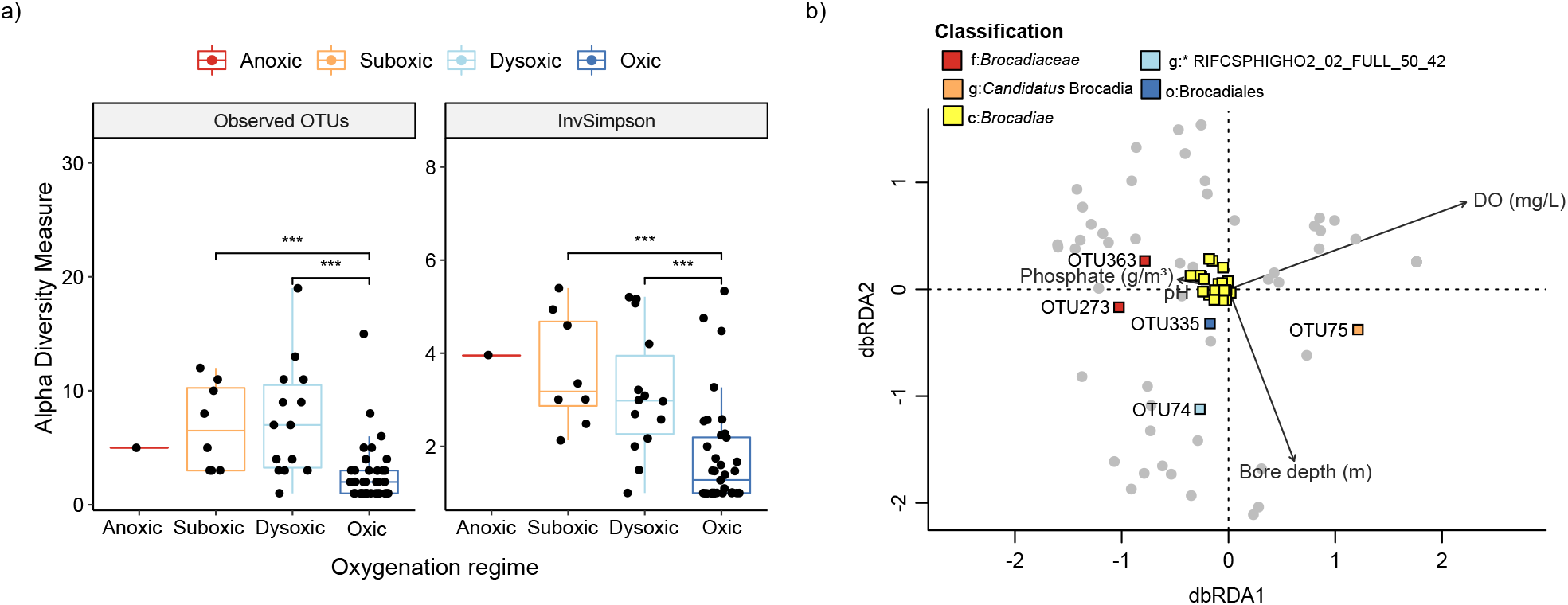
Diversity analysis and an ordination plot showing the environmental variables and factors influencing the structure of the anammox community. **a)** Boxplots represent observed OTUs (richness) and Inverse-Simpson diversity at anoxic, suboxic, dysoxic and oxic sites (Significance levels = *p* < 0.01; Wilcoxon Test). **b)** Distance-based Redundancy Analysis of the 16S amplicon data from the 60 groundwater samples using a Bray-Curtis dissimilarity matrix between samples based on OTU abundance. Vectors show significant environmental variables (excluding ORP due to missing values), constraining the variability in community composition. *Brocadiae* OTUs were added to the ordination using species scores (shown as large coloured squares) and the 5 OTUs shown are identified to the lowest level of taxonomy (Acronyms: g = genus; f = family; o = order; c = class; p = phylum).

Oxygen tolerance in anammox bacteria has been demonstrated previously [77,78], particularly for *Ca*. Brocadia. For example, *Ca*. Brocadia sinica exhibited tolerance in the presence of >1 mg/L DO *in vitro* [79], and anammox bacteria closely related to *Ca*. Brocadia fulgida have been reported in groundwater with up to 3 mg/L oxygen [8] (>3 mg/L DO is generally classified as oxic [29]). A recent study analysing the metabolism of *Ca*. Brocadia species, identified genes encoding superoxide dismutase and cytochrome *c* peroxidase genes that are involved in oxygen detoxification [80], as may be expected for organisms living at reduction-oxidation interfaces. Although *Brocadiae* overall and *Ca*. Brocadia were detected in groundwater with much higher DO concentrations in this study (up to 10.6 mg/L), aquifers are spatial heterogenous, and *Brocadiae* likely inhabit lower oxygen niches within the sampled aquifers. In comparison, a further 25.9% of *Brocadiae* comprised *Planctomycetes* RIFCSPHIGHO2_02_FULL_50_42 (mostly OTU 75). The overall relative abundance was significantly and negatively correlated with DO (*r=*-0.38), ORP (*r*=-0.45) and NO_3_^-^ (*r=*-0.29), indicating a preference for reducing conditions (Table S8). It was originally recovered from an aquifer adjacent to the Colorado River (Rifle, USA) [56], with conditions similar to site (Wel13) here, where the genus was most abundant, including identical dysoxic conditions (0.78 mg/L DO) with low NO_2_^-^ and NO_3_^-^.

#### DOC did not negatively impact Brocadiae community abundance

Anammox generally occurs at low organic carbon levels [81]. As reported above, DOC and *Brocadiae* community abundance were instead positively correlated. This association may be best explained by the negative relationship typically shared by DO and DOC in groundwater, where organic carbon availability regulates aerobic metabolism [82], although DO and DOC were not significantly correlated in this study (*r*=-0.08, *p*=0.5), possibly due to DOC being rapidly utilized in groundwater. Nevertheless, some anammox bacteria (e.g. *Ca*. Anammoxoglobus propionicus) have a better affinity for NO_2_^-^ in the presence of organic acids, such as propionate, or can assimilate formate via the Wood–Ljungdahl pathway (as recently shown for *Ca*. Kuenenia stuttgartiensis) [83], implying that groundwater DOC could support the growth of some anammox bacteria.

#### Hydrazine synthase capacity, but not activity, was correlated with DO

To confirm anammox potential and activity in groundwater, we quantified genes and transcripts encoding hydrazine synthase subunit *hzsB*. Together, genes *hzsA, hzsB* and *hzsC* encode hydrazine synthase, which is a key enzyme in anammox metabolism, converting nitric oxide and ammonium into hydrazine [6]. The relative abundance of *Brocadiae* 16S rRNA gene sequences, correlated strongly with *hzsB* gene copies (*r*=0.62) (Fig. 3a), consistent with the expectation that *Brocadiae* undertake anammox [84]. The concentration of *hzsB* genes ranged from 3.64 × 10^1^ to 1.23 × 10^7^ genes/L of groundwater (excluding 11.1% of samples below detection) with an average of 4.9 × 10^5^ ± 2.1 × 10^6^ SD genes/L. This is consistent with concentrations of ∼1.5 × 10^4^ to ∼1.0 × 10^8^ genes/L found elsewhere in the subsurface, including in soil horizons interacting with the water table [8,85]. The average number of *hzsB* transcript copies was significantly lower than for *hzsB* gene copies (Wilcoxon test *p* <0.01; 4.5 × 10^5^ copies/L less, ∼13.2 times lower) (Fig. 3b), suggesting a latent capacity of 92.4%. Overall, *hzsB* gene copy numbers were negatively correlated with bore depth (*r*=-0.34) and DO (*r*=-0.42), reflecting the trend seen with *Brocadiae* 16S rRNA gene relative abundance and DO, and as expected for an anaerobic process. However, no significant relationship was evident between *hzsB* gene transcripts or anammox activity and oxygen availability (*r*=-0.16, *p*=0.44) or bore depth (*r*=0.14, *p*=0.5).

**Figure 3.**
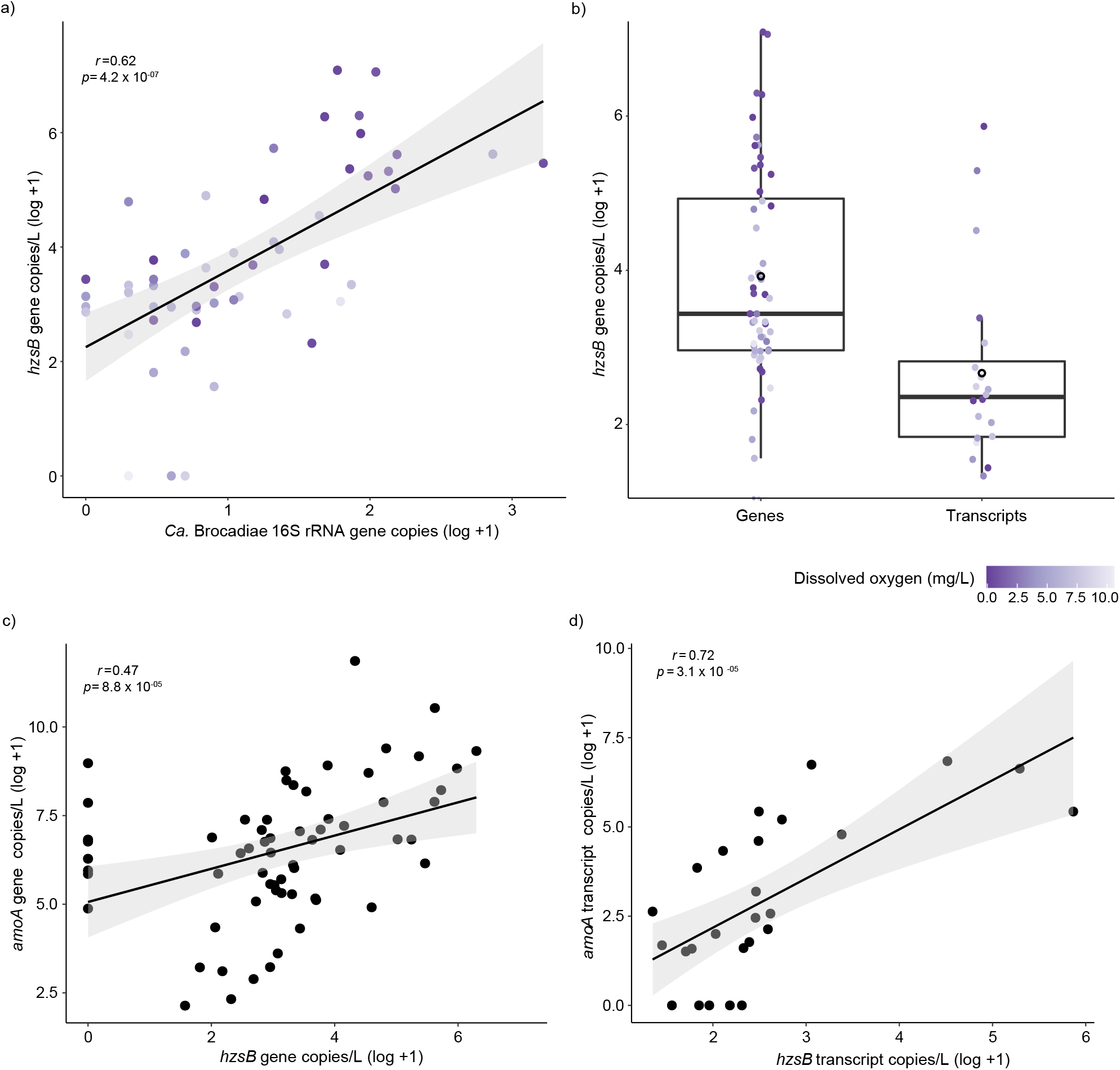
Correlations between 16S rRNA gene amplicon data, aerobic ammonia oxidisers (AOA and AOB) and abundance of hydrazine synthase genes and transcripts. **a)** Correlation of *hzsB* gene per L of groundwater (log10) and number of *Brocadiae* (log10) sequences within the rarefied OTU table with samples containing class *Brocadiae*. **b)** Boxplot representing log10 abundance of *hzsB* genes for genes and transcripts. Black open circle represents the mean and horizontal line represents the median of gene copies per L. Purple gradient for **a)** and **b)** correspond with DO concentration (mg/L). **(c)** Correlation of *amoA* genes (log10) and *hzsB* genes (log10) copies per L of groundwater using Spearman correlation for genetic potential (DNA). **d)** Correlation of *amoA* genes (log10) and *hzsB* genes (log10) copies per L of groundwater using Spearman correlation in transcripts (RNA).

### Co-occurrence of aerobic and anaerobic ammonia oxidizers

The nitrification-anammox process allows for elevated N-removal efficiencies from NH_4_^+^ in naturally ammonium-rich or contaminated water [86], by ensuring a portion of ammonium is converted to nitrite. Potential combined nitrification-anammox was determined by quantification of archaeal and bacterial ammonia monooxygenase genes (*amoA*). Results show *hzsB* and archaeal and bacterial *amoA* genes (*r*=0.47) and transcripts (*r*=0.72) were strongly correlated, suggesting co-occurrence of aerobic and anaerobic ammonium oxidation (Fig. 3c-d). Co-occurrence has been observed previously within lab-based models [87], and an aquifer system, where anammox was the dominant process [8]. Together, these results suggest that anammox in aquifers relies, at least partially, on nitrite derived from aerobic ammonia oxidation. Results here also indicate anammox is more frequently the dominant process when considering a wide range of aquifers. Aerobic ammonia monooxygenase gene copies were more abundant in 83% of samples compared to hydrazine synthase genes; however, hydrazine synthase transcripts were higher in 69% of samples.

### Phylogenomic diversity of anammox bacteria in aquifers

Genomes were reconstructed from 16 samples (gwj01-gwj16) across 8 wells (groundwater ± biomass-enrichment) spanning a range of oxygen (0.37–7.5 mg/L) and nitrate (0.47–12.6 g/m^3^) concentrations (Table S1). Of 541 MAGs (>50% complete, ≤5% contamination), 8 were identified as *Ca*. Brocadiae (63.4–95.6% complete, Table S10). Of these MAGs that contained partial or (near)full length (445-1496 bp) 16S rRNA gene sequences (nzgw513, nzgw516-517), those from nzgw516-517 aligned fully with OTU729 (272 bp long), which was present in 21/80 samples (total of 92 sequences). Both the nzgw516-517 and OTU729 sequences had best hits to *Planctomycetes* spp., including *Planctomycetes* bacterium strain Pla86 and *Ca*. Kuenenia stuttgartiensis when searched against the NCBI NR database (BLASTN; 79.54%-82.85% identity), consistent with phylogenomic analyses showing these genomes comprise a novel clade (Clade II, Fig. 4a). OTU729 was only classified to the domain Bacteria using USEARCH with the SILVA SSU Ref NR99 database, and RDP Classifier, suggesting that these novel *Ca*. Brocadiae could be underestimated in the environment when using a 16S rRNA gene based approach.

**Figure 4.**
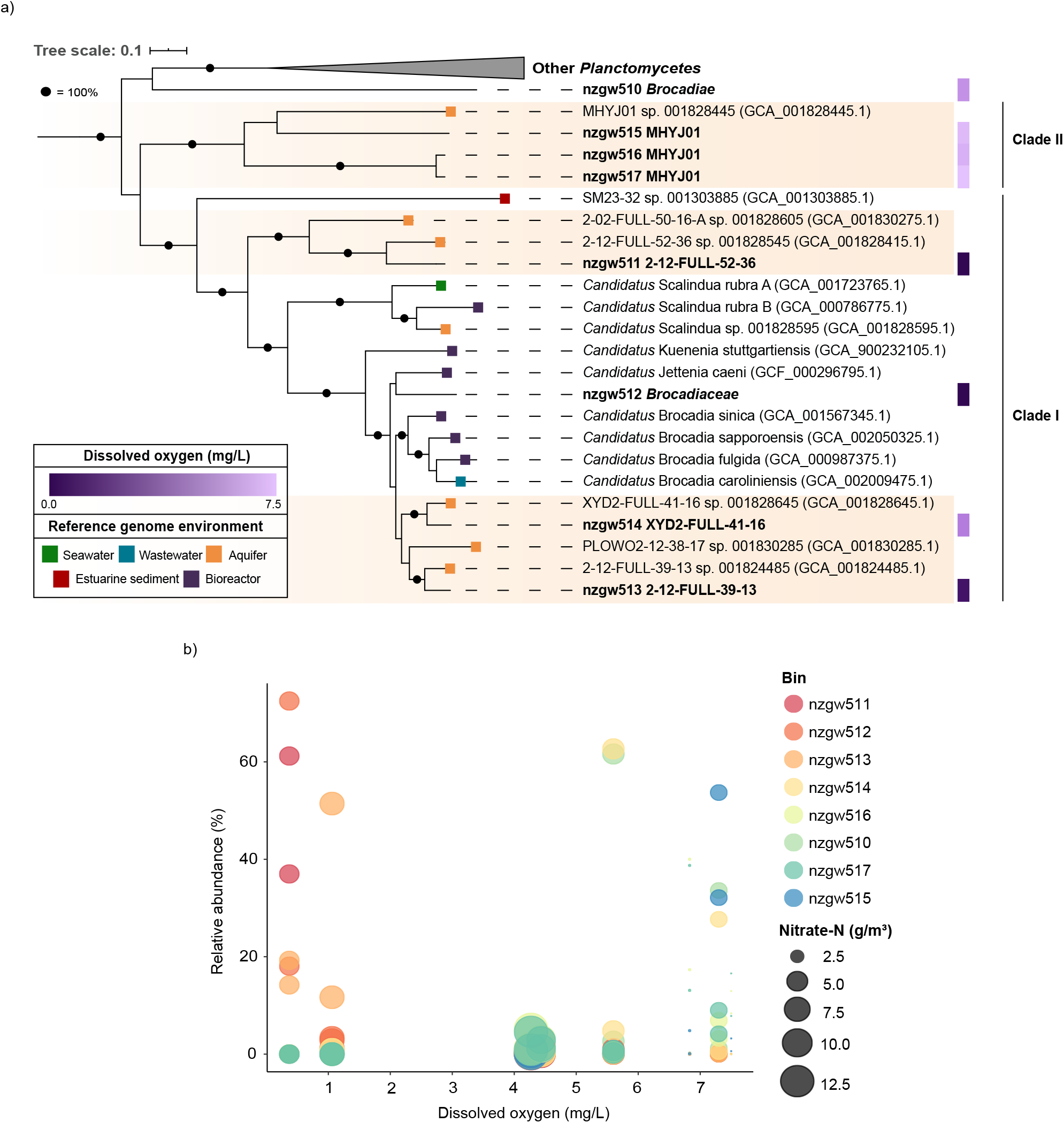
Phylogenetic distribution and relative abundance of *Brocadiae* MAGs. **a)** Maximum-likelihood phylogenomic tree of 28 *Planctomycetes* genomes based on 120 concatenated bacterial marker genes (GTDB-Tk) with 5,040 amino-acid sites using LG+F+R5 model of substitution and 1,000 bootstraps. Purple gradient represents the DO concentration (mg/L) at the site with highest relative genome abundance for that genome. Coloured tiles represent the environment of recovery for the reference genomes. Bootstrap values shown as black circles equal 100%. Scale bar indicates the number of substitutions per site. Sequences from this study are shown in bold font, with both the study identifier and GTDB classification given. Orange colour strip represents clades of *Brocadiae* genomes recently recovered from aquifers. See Table S5 for reference genome details. **b)** *Brocadiae* genome relative abundance across DO concentration (mg/L) at each site (relative to other *Brocadiae*). Bubble size corresponds to nitrate concentration (g/m^3^) at each site.

A pairwise comparison with 16 other *Brocadiae* genomes from various environments yielded ANI values below the proposed species cut off of 95% [88] (Fig. S5), implying the newly recovered MAGs represent novel species. The phylogenomic tree (Fig. 4) revealed six NZ aquifer-derived MAGs form three (sub-)clades with recently recovered genomes from the aquifer in Rifle, Colorado [56], suggesting these are groundwater adapted species. One (nzgw512) clustered with well-characterised anammox bacterium, *Ca*. Jettenia caeni [89]. An eighth MAG, nzgw510, was phylogenetically distinct from other *Ca*. Brocadiae. MAGs clustering with Rifle MAG “MHYJ01” (from aquifer sediment [56]) constitute Clade-II, which is distinct from well-known anammox bacteria (Clade-I, Fig. 4a). They contained a much higher GC content (GC >67%) than others from this study, excluding outlier nzgw510 (average 42.5% ± 3 SD) (Table S10). Higher GC may indicate a higher evolutionary rate for these organisms [90]. All lower GC *Brocadiae* MAGs from this study, including two aquifer subclades, co-clustered with well-known anammox *Candidatus* genera from enrichment studies (Clade-I). The estimated genome sizes were significantly smaller than the high GC group (Clade-I: 3.22 Mbp ± 0.92, Clade-II: 5.16 Mbp ± 1.24), further suggesting a divergence in evolutionary strategy employed by the two groups.

As for 16S rRNA gene analyses (Fig. 1b), *Brocadiae* MAG relative abundances varied across sites, corresponding to distinct geochemical conditions (Fig.4a, Table S10) but not strictly phylogenetic relatedness (Fig. 4b). Three of four MAGs associated with well-characterized Clade I anammox-bacteria (nzgw511-513) were most abundant at sites with low DO (<1.1 mg/L) and high DOC (3–26 g/m^3^). MAG nzgw514 is one of two closely related to *Ca*. Brocadia. Consistent with 16S rRNA gene data (Fig. 1b) it demonstrated a preference for oxic groundwater (Fig. 4b). Together with the high-GC MAGs (nzgw510 and Clade II), it was more abundant at sites with high DO (5.6–7.5 mg/L) and low DOC (0–0.8 g/m^3^). Factors identified that influence ecological niche differentiation among anammox species, include microbial growth kinetics, organic matter, oxygen tolerance, aggregation ability and interspecific competition [5]. Collectively, results here suggest DO availability may be a major driver of anammox bacterial diversification in groundwater.

### Genomic and transcriptomic evidence for anammox metabolism and oxygen tolerance

#### Characteristic hydrazine based steps in anammox

Nitric oxide produced by NO_2_^-^ reductase [91], or by cyclic feeding, is used by hydrazine synthase (HZS), along with NH_4_^+^, to produce hydrazine (N_2_H_4_) within the anammoxosome [6]. Most *Brocadiae* MAGs recovered (6 of 8) have genes encoding signature anammox steps for hydrazine production (hydrazine synthase) and removal (hydrazine dehydrogenase) (Fig. 6; Table S11) [91]. This excludes phylogenetic outlier nzgw510 and Clade-II nzgw517, with the lowest genome completeness (75% and 63%, respectively). Nonetheless, HZS genes in nzgw515-516 indicate that Clade-II bacteria are capable of anammox [91], and may be important contributors to anammox in oxygen-rich groundwaters (Fig. 4b) [8]. The final anammox step is catalysed by an octahaem-hydrazine dehydrogenase (HDH/*hzoA*). N_2_H_4_ is oxidised using cytochrome *c* as an electron acceptor [92]. Although no Clade-II MAGs contained *hzoA*, nzgw515 contained hydroxylamine oxidoreductase, which is also capable of hydrazine oxidation *in vitro* [93]. Multiple *hzoA* gene copies are present in nzgw511–514, consistent with close phylogenetic relatedness to well-characterised anammox bacteria (Fig. 4).

Reinforcing genomic evidence, metatranscriptomics data for six samples (gwj09, gwj11, gwj13-16) across two sites, revealed that the transcriptional activity of *hzoA* in MAGs nzgw511-514. This activity was 356-fold higher at the dysoxic site (wells E1 and N3) than the oxic site (Fig. 6b), and likely contributed in part to excess N_2_ measured at those sites (Fig. 5). Despite the compositional bias in anammox bacteria associated with differences in DO (Fig. 4), results confirm that in groundwater with low oxygen concentrations anammox bacteria are more active [8]. Contemporaneous measurement of excess N_2_ indicated active N_2_ generation in dysoxic groundwater (from wells E1 and N3) due to denitrification or anammox. In contrast, oxic groundwater was either at the bounds of uncertainty (wells BW8 and RF3) or devoid of measurable excess N2 (from wells SR1-2, BW19, RF2) (Fig. 5), which, accordingly, included groundwater with relatively little observed hydrazine synthase/dehydrogenase activity (from wells SR1-2). Additional quantification via ddPCR of hydrazine synthase transcripts from each site, sampled at an earlier date, demonstrated that anammox was active in groundwater associated with all *Ca*. Brocadiae MAGs, which were present in both dysoxic to oxic sites (Fig. 4b) distributed across 8 wells over 4 sites (Table S9). However, quantitative results were consistent with relative metatranscriptomics data, indicating transcriptional activity was 100-fold higher in dysoxic versus oxic groundwater in the four samples analyzed (Fig. S6).

**Figure 5.**
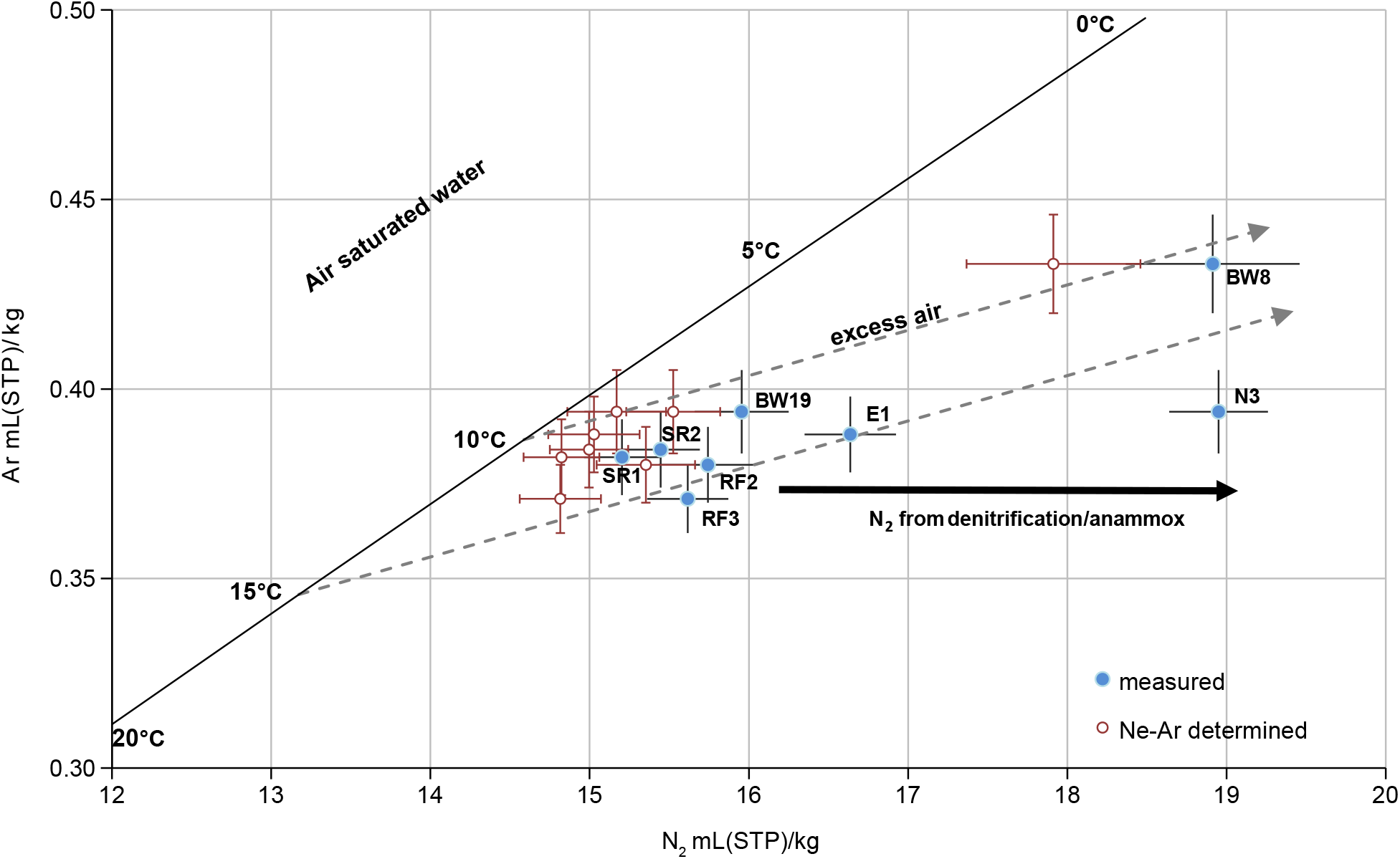
Plot of dissolved nitrogen versus dissolved argon concentrations at sites sampled for metagenomics analysis. Dissolved Ar, N_2_ are expressed in mL of the respective gas at standard temperature and pressure (STP=273.15 K, 101.325 kPa) per kg of water. Bold line represents gas concentrations in water which are in equilibrium with the atmosphere at the given temperature. Arrows indicate competing processes that can alter gas concentrations, where grey dashed lines indicate excess air in groundwater relative to atmosphere (with upper and lower lines representing addition of unfractionated excess air relative to equilibrium concentrations at 10 and 15°C). Black horizontal arrow depicts additional excess N_2_ inferred to be from biological processes (denitrification or anammox). Reconstructed N_2_ data (in equilibrium with inert atmospheric gases), based on groundwater recharge temperatures and excess air concentrations derived from dissolved Ne and Ar data (shown as red hollow symbols). Recharge temperature = temperature of recharging water at the time it enters the groundwater system. Excess air = dissolved air in excess of the equilibrium soluble amount at the given recharge temperature (thought to originate from processes such as bubble entrapment occurring during recharge and subsequent dissolution under increased hydrostatic pressure). The difference between these and the measured N_2_ data (blue solid symbols, and shift their along x-axis relative to red hollow symbols) indicates the amount of N_2_ in excess, formed via denitrification and/or anammox at each site. Error bars show the combined statistical standard uncertainty from all processes and calculations contributing to the measurement uncertainty, expressed as one standard deviation. Groundwater from wells SR1-2, BW8, BW19, RF2-3 is characterised as oxic, while groundwater from E1 and N3 is dysoxic-suboxic (Table S1).

**Figure 6.**
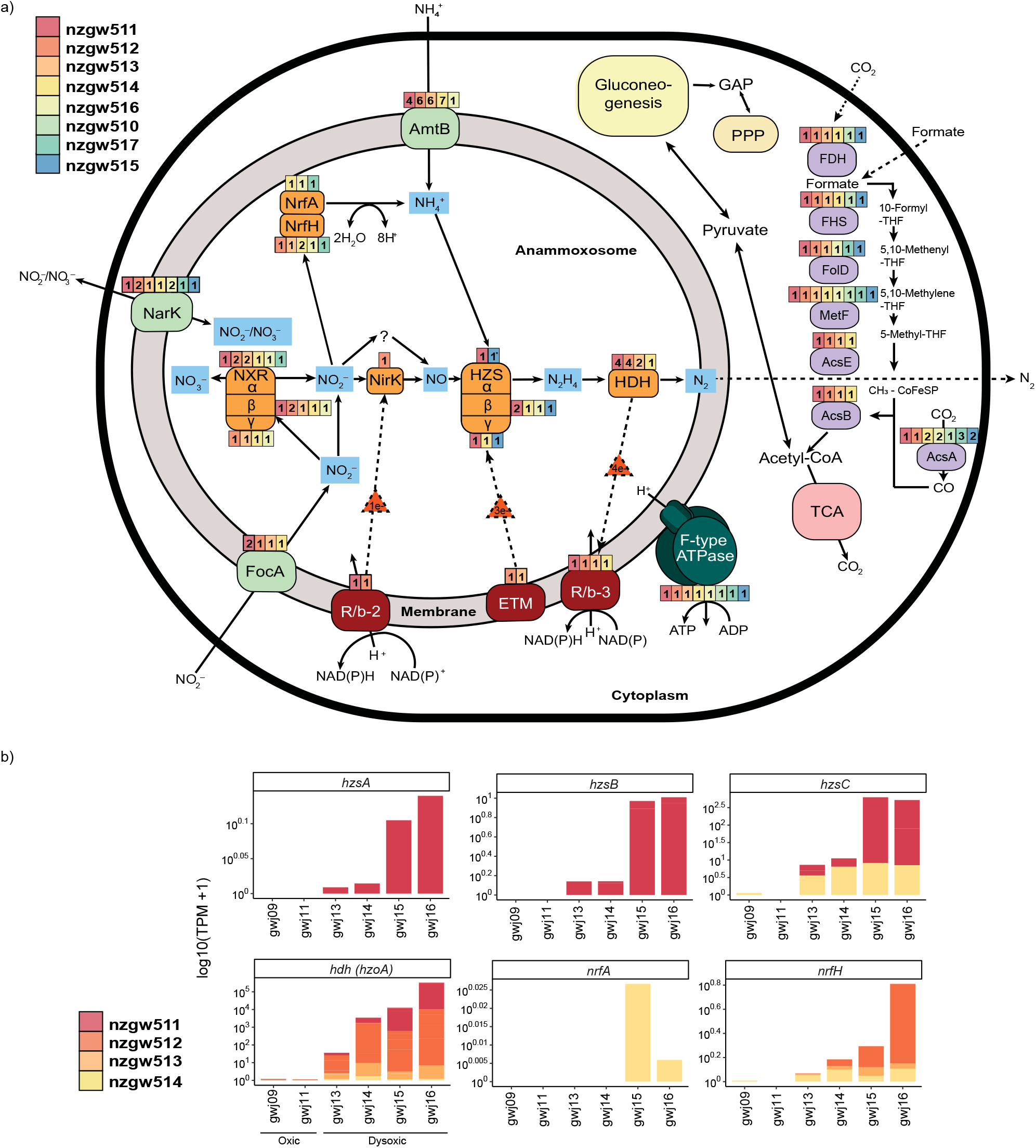
An overview of the predicted metabolic pathways of characterised anammox bacteria built from the summarised annotated data from the reconstructed genomes and mapped transcripts. **a)** Metabolic pathways present in MAGs. Nxr, nitrite:nitrate oxidoreductase; Nir, nitrite reductase; Nrf, nitrite reductase forming ammonium; HZS, hydrazine synthase; HDH, hydrazine dehydrogenase; AmtB, ammonium transporters; FocA, nitrite transporters; NarK, nitrite/nitrate transporter; ETM, electron transfer module from quinone pool to HZS(composed of kuste2856 and kuste2855); R/*b*, Rieske/cytochrome b (bc_1_) complexes, R/*b*-2 (kustd1480-85), and R/*b*-3 (kuste4569-74); F-type ATPase, F-type ATP synthase (MAGs containing ≥50% of subunits); GAP, Glyceraldehyde 3-phosphate; EMP, Embden-Meyerhof-Parnas pathway; FDH, Formate dehydrogenase; FHS, Formate--tetrahydrofolate ligase; FolD, methylenetetrahydrofolate dehydrogenase; MetF, Methylenetetrahydrofolate reductase; AcsE, 5-methyltetrahydrofolate:corrinoid; AcsB, acetyl-CoA synthase; AcsA acetyl-CoA synthetase; PPP, Pentose phosphate pathway; TCA, tricarboxylic acid cycle. Colours represent genomes from this study, number equals copies present.* = Partial HzsA subunit. **b)** Logged (log 10) TPM values for active hydrazine synthase, hydrazine dehydrogenase and nitrite reductase forming ammonium genes from groundwater characterised as oxic (samples gwj09 from well SR1 and gwj11 from well SR2) and dysoxic (gwj13-14 from well E1 and gwj15-16 from well N3). Samples gwj14, gwj16 are of sonicated groundwater.

Further analyses showed HZS sequence similarity and structure did not reflect *Brocadiae* phylogenomic relatedness. All three HZS subunits encoded by nzgw511 are closely related to *Ca*. Kuenenia stuttgartiensis subunits (Fig. 7). However, HzsB from Clade-II nzgw516, and Clade-I nzgw514, and nzgw511 (duplicate), constitute a distinct HzsB clade relative to other anammox bacteria, indicating evolutionary divergence. Clade-II nzgw515 instead encodes for a fused HZS-beta-gamma protein subunit, comparable to distant *Brocadiae* relatives *Ca*. Scalindua profunda [94] and Scalindua brodae [95].

**Figure 7.**
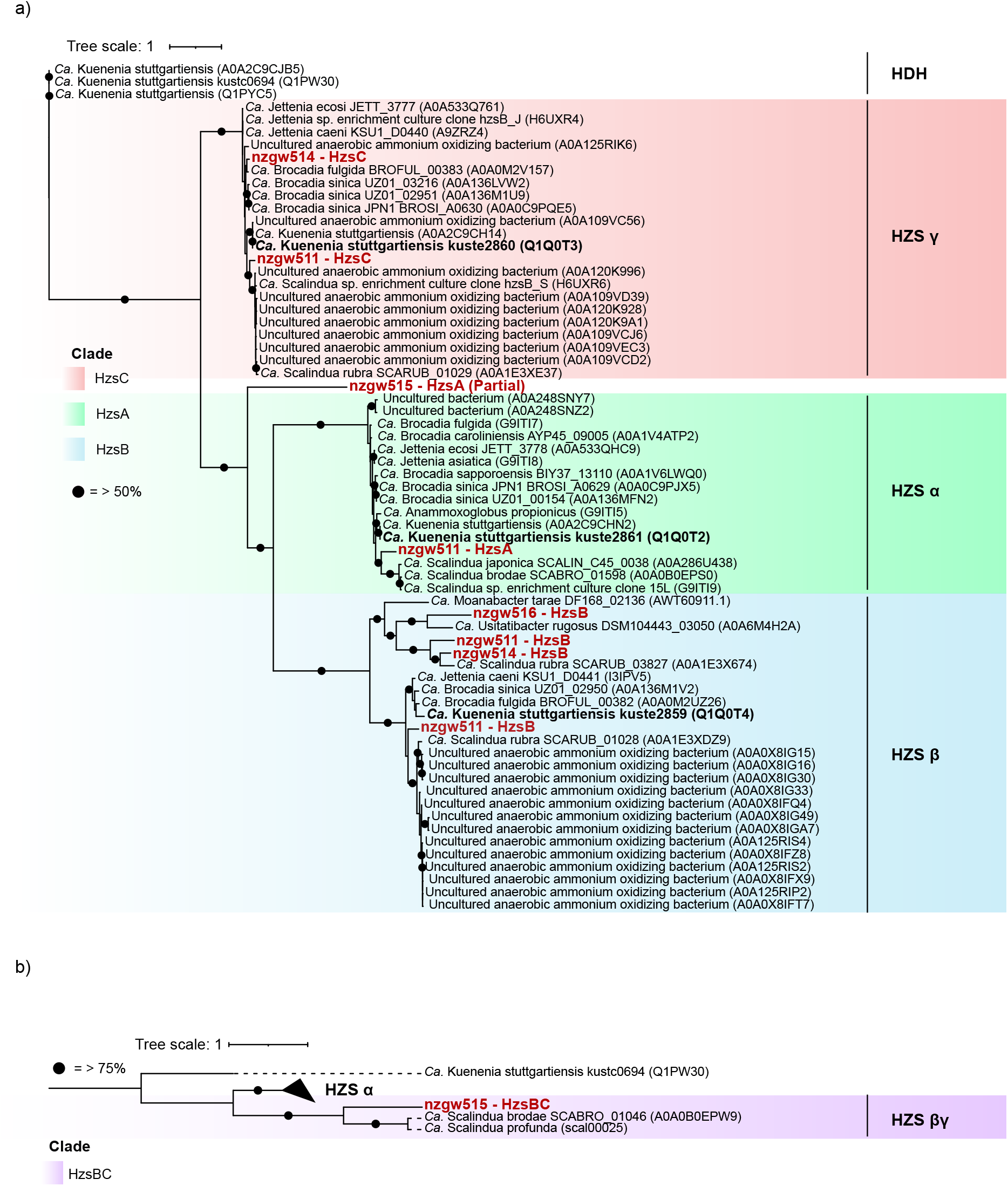
Phylogenetic tree of the recovered hydrazine synthase protein subunits from *Brocadiae*. **a)** Phylogenetic tree with 65 protein sequences of hydrazine synthase subunits with 964 amino-acid sites, and built using WAG+G4 model using 1,000 bootstraps. The hydrazine synthase (HZS) alpha, beta and gamma proteins were predicted from protein coding sequences recovered from genomes in the study (red) and other HZS protein sequences available from Uniprot database. HDH = Hydrazine dehydrogenase. Bold black sequences are from representative model organism *Ca*. Kuenenia stuttgartiensis. Circles represent > 50% bootstrap value. Scale bar represents number of substitutions per site. **b)** Phylogenetic tree of fused Hzs beta-gamma proteins (from predicted protein coding sequences) made with 10 sequences (6 HzsA subunits) and 903 amino-acid sites, and built using WAG+G4 model (1,000 bootstraps). Circles represent > 75% bootstrap value, red sequence was recovered in this study.

#### Production of nitric oxide (and ammonium) for anammox

In the typical anammox reaction, the first step is NO_2_^−^ reduction to NO. Nitrite, in addition to other external substrates, such as nitrate, nitrite and ammonium, are acquired by transporters (see Supplementary Information for details on transporters in MAGs). NO_2_^−^ reduction to NO is usually catalysed by cytochrome *cd*_1_-type nitrite reductase (*nirS*), as found in *Ca*. Kuenenia and *Ca*. Scalindua [94,96], or the copper-containing nitrite reductase (*nirK*), as for *Ca*. Jettenia spp. [97]. One MAG (nzgw512) contained *nirK* similar to *Ca*. Jettenia caeni [89]; however, other MAGs were devoid of *nirK* and *nirS* (Fig. 6). Canonical NO-forming NO_2_^−^ reductases are also lacking in several *Brocadia* species (e.g. *Ca*. Brocadia sapporoensis and *Ca*. Brocadia sinica) [98] and Rifle aquifer genomes (e.g. 2-12-FULL-39-13 and 2-02-FULL-50-16-A) [56]. Some anammox bacteria are proposed to utilize different enzymes to reduce NO_2_^−^ to NO [99]. These enzymes may be among the hydroxylamine oxidoreductase like octaheme proteins encoded by anammox genomes [91]. Within *Ca*. Kuenenia stuttgartiensis, this protein (kust0457-kust0458), comprising octaheme kustc0458 and diheme kustc0457, constitutes a heterododecameric (α_6_β_6_) complex comprising 60 *c*-type hemes [91]. The second is kuste4574, a close homolog of kustc0458, which forms part of a novel Rieske– heme *b* (R/*b*; *bc*_1_) complex (kuste4569-kuste4574). It is postulated that these proteins catalyse NO_2_^-^ reduction to NO [91]. Protein-coding sequences similar to kustc0457-kustc0458 and kuste4574 were identified in nzgw511, nzgw513-514 (Table S6). These proteins may represent an alternative pathway for reducing NO_2_^-^. Furthermore, *Ca*. Kuenenia stuttgartiensis can grow in the absence of NO_2_^-^ by coupling ammonium oxidation to NO reduction, producing N_2_ without N_2_O emissions [100]. This suggests that anammox bacteria may mitigate natural NO and N_2_O emissions in many ecosystems.

It is suggested that anammox bacteria adapt a Dissimilatory Nitrate Reduction to Ammonia (DNRA) like metabolism in the absence of NH_4_^+^, using nitrate reductase, followed by NrfAH cytochrome *c* nitrite reductase [72]. The respective products, NO_2_^-^ and NH_4_^+^, can then drive anammox. We found *nrfAH* genes in phylogenetically diverse genomes (nzgw514, nzgw516-517). Further, genomes nzgw512–514 contain a tetraheme cyctochrome *c nrfH* gene, with CxxCH repeated sequence motifs similar to *Ca*. Brocadia [99] and *Ca*. Jettenia [97]. Both *nrfA* and *nrfH* genes were observed to be expressed by these taxa in dysoxic groundwater (Fig. 6b). Together, these genes and transcripts indicate the active reduction of NO_2_^-^ to NH_4_^+^ in a NAD(P)H-dependent, six-electron transfer reaction, via a mechanism resembling DNRA [91], potentially fuelling active anammox in the same sites.

NAD(P)H nitrite reductase genes (*nirD* and/or *nirB*), which also reduce NO_2_^-^ to NH_4_^+^, were found in Clade-II genomes, nzgw515–517. The large subunit encodes NirB, an iron–sulfur protein. All protein-coding sequences had the highest identity (79–84%) to NirB from *Planctomycetes* genome (GCA_016200125.1), recently recovered from a pristine aquifer in the USA [101]. This had a similar geochemical profile to the site where nzgw516-517 were most abundant (low NH_4_^+^, <0.05 mg/L, and NO_3_^-^, 0.23 mg/L). The *nirB* gene was recently also identified in a marine-derived anammox bacterium, related to *Ca*. Scalindua [102]. We also identified a small subunit, NirD, in nzgw516-517. The sequence contains a Rieske non-heme iron oxygenase family domain with closest similarity to *Verrucomicobia* species (63.5%). NH_4_^+^ often limiting in groundwater [20]. In this study, NH_4_^+^ was undetectable in groundwater from which MAGs were recovered, and detected at only 31% of sites overall. The presence of these Nrf and Nir proteins could therefore mediate an important alternative process providing NH_4_^+^for anammox [91].

#### Mechanisms of oxygen tolerance in groundwater anammox bacteria

All *Brocadiae* genomes encode proteins for oxygen protection, such as BatD and BatA, which are involved in oxygen tolerance in *Bacteroides* species [103], alkyl hydroperoxide reductases and thioredoxin [104]. Thioredoxin and peroxiredoxin genes were expressed in MAG nzgw510 at the oxic site (0.06-0.15 TPM), and BatD, BatA, catalase, peroxiredoxin and thioredoxin were expressed by MAGs nzgw511-514 at the dysoxic site (0.112-2.81 TPM). Proteins encoding respiratory chain cytochrome *c* oxidase, cbb3-type (ccoN, ccoO and ccoP) were all or partially present in four MAGs (nzgw511-14). Cbb3 is a multi-chain transmembrane protein located near the anammoxosome membrane [105]. This enzyme is believed to have evolved to perform a specialized function in microaerobic energy metabolism, however it could also merely act in oxygen detoxification [72]. Additionally, three Clade-II genomes encode a caa3-type cytochrome oxidase, containing unique CoxA/B subunits. In *Acidithiobacillus ferrooxidans coxA/B* genes encode the terminal part of the ferrous iron “downhill” pathway, which shuttles electrons from extracellular iron to oxygen [106], and thus represents another potential detoxification strategy.

#### Alternative metabolic pathways in groundwater anammox bacteria

Anammox grow chemolithoautotrophically by performing CO_2_ fixation via the Wood-Ljungdahl pathway [107]. The products of this reaction, acetyl-CoA and pyruvate, enter the oxidative TCA cycle and gluconeogenesis (Fig. 6; Table S10; see details on TCA cycle in Supplementary Information). Several ABC-transport systems were present within the genomes, revealing variations in substrate importation such as phosphate, cobalt, nickel, iron (III), zinc, sulfate, molybdate, lipoproteins, ribose, rhamnose, polysaccharides and oligopeptides (Table S10). Oligopeptide transport systems were present in three genomes (nzgw515-517), as seen in *Ca*. Scalindua profunda, and suggest these genomes are capable of oxidizing decaying organic matter [108], in addition to carbon fixation via the Wood-Ljungdahl pathway (Fig. 6). Additionally, iron (III) transporters were present in three genomes which may enable the coupling of formate oxidation to iron (III) reduction in the absence of NO_2_^-^ or NO, as previously identified in other species [109]. Moreover, hydrogenase genes were also present in three of the genomes. The first encodes a [Ni-Fe] group 3b hydrogenase, found in nzgw511, which is proposed to harbour (sulf)hydrogenase activity [110]. This genome also encoded a sulfate adenylytransferase, which may play a role in sulfide oxidation [111]. The second is a group 4 hydrogenase present in nzgw513. These hydrogenases encode coupled oxidation of NADPH to the evolution of H_2_ [112], indicating hydrogen turnover and/or an alternative energy source. Our results therefore suggest these groundwater anammox bacteria are metabolically versatile, containing various hydrogenases and ABC transporters for organic compounds, and genes encoding DNRA, which would allow for growth in substrate limiting conditions (such as low ammonium concentrations).

## Conclusions

This study shows that anammox bacteria are prevalent and active across wide-ranging aquifer chemistries and lithologies, including in oxic groundwater, although predicted to be unfavourable. While we found significantly more *Brocadiae* diversity in anoxic groundwater, some taxa were positively associated with DO concentrations. Of the eight novel *Brocadiae* genomes reconstructed, three belong to under-characterised clades only previously recovered from aquifers. Most possess genes involved in signature pathways of anammox, including novel hydrazine synthase genes. A co-occurrence of anaerobic and aerobic ammonia-oxidisers at many sites suggests metabolic hand offs (such as nitrite) between these processes. Furthermore, genomic characterization of *Brocadiae* identified a range of potential aero-tolerance mechanisms, explaining our finding of anammox in oxic groundwaters. Results indicate niche differentiation amongst anammox bacteria based on oxygen concentrations, and that anammox is a common mechanism for nitrogen removal in aquifers.

## Declarations

### Ethics approval and consent to participate

Not applicable.

### Consent for publication

Not applicable.

### Availability of data and materials

All amplicon, WGS and metatranscriptome data from groundwater and biomass-enriched groundwater samples have been deposited with NCBI under BioProject PRJNA699054.

### Authors’ information

#### Authors’ contributions

OM, EG and KMH collected samples. OM and EG led field trips. OM, EG, KMH, LW and MC planned the study. OM and EG conducted the laboratory work. OM and EG analyzed and interpreted data. RR and HM quantified excess N_2_ gas at four sites. OM wrote the first draft of the manuscript. OM, KMH and EG revised and corrected the manuscript. All authors read and approved the final manuscript.

#### Corresponding author

Correspondence to Kim M. Handley.

## Acknowledgements

We thank P Abraham (ESR), HS Tee and JS Boey (University of Auckland) for help sampling Canterbury wells, and council staff in Auckland (C Foster), Wellington (D McQueen), Waikato (S Herath) and Canterbury (D Evans and R Cressy) for help accessing and sampling wells. We thank D Waite and JS Boey for bioinformatics support. Computational resources were provided by New Zealand eScience Infrastructure.

