## Supplementary Information for "Metabolic diversity and aero-tolerance in anammox bacteria from geochemically distinct aquifers"

**Supplementary Methods**

*Chemical analysis of water samples*

Total phosphorus and phosphate were determined according to American Public Health Association (APHA) 4500-P B & E (modified to include an acidic ammonium persulphate to convert organophosphates and polyphosphates to orthophosphate) [1]. Dissolved reactive phosphorus (DRP) was determined according to APHA 4500-P G (sample was reacted with ammonium molybdate and ascorbic acid to form molybdenum blue then detected at 880 nm). Total ammoniacal-N was determined according to APHA 4500-NH3 H (using phenol/hypochlorite reaction forming a complex that was detected at 630 nm) and calculated as NH4-N = NH_4_^+^-N + NH_3_-N. Nitrite-N and Nitrate-N + Nitrite-N were determined according to APHA 4500-NO_3_ I (NOxN is measured via automated cadmium reduction and griess reaction. NO_2_N calculated by automated griess reaction (Sulfanilamide) detected at 540nm). Nitrate-N was calculated by (Nitrate-N + Nitrite-N) – Nitrite-N. Total organic carbon (TOC) and dissolved organic carbon (DOC) were measured according to APHA 5310 C (Analysed using Super Critical Persulphate Oxidation with phosphoric acid and sodium persulphate), where TOC = total carbon – total inorganic carbon. For DOC, groundwater was filtered first using 0.45 μm Polypropylene filter. Total suspended solids were measured by first evaporating groundwater samples in an oven at 105 °C until dry. Dried solids were then weighed and normalized to the total water volume analyzed. Sulfate was measured according to APHA 4110 B. Alkalinity and dissolved manganese were analysed according to APHA 2320 B and APHA 3125 B, respectively.

**Supplementary Results and Discussion**

*Mechanisms for nitrate, nitrite and ammonium uptake and nitrate reduction*

To fuel anammox, the externally acquired substrates, NH_4_^+^ and NO_2_^−^, need to cross both the cytoplasmic and anammoxosome membranes [86]. Multiple copies of NH_4_^+^, NO_2_^−^ and NO_3_^−^ transporters that could facilitate transport into the anammoxosome are found in all published anammox genomes [79, 84, 85, 87, 88]. Five of our newly recovered genomes possess one to seven copies of *amtB*-like ammonium-transporter genes (Fig. 6), which neighboured genes encoding the nitrogen-regulatory protein P-II. This protein binds directly to AmtB and regulates the ammonia channel [89]. Genes encoding putative NarK NO_3_^−^/NO_2_^−^ transporters were present in seven MAGs (1—2 copies each). Nitrite transport is also mediated by a bidirectional NO_2_^−^/formate transporter gene, *FocA* [62], which is present in four of the groundwater MAGs (1—4 copies each). Finally, interconversion of NO_3_^-^ and NO_2_^-^ is catalysed by a nitrate reductase (NXR nitrate:nitrite oxidoreductase) [62]. NXR genes were present in six genomes.

*TCA cycle in anammox bacteria*

*Ca.* Kuenenia stuttgartiensis does not encode a citrate synthase which is typically required to operate the oxidative TCA cycle, and here, it was absent in five of the reconstructed groundwater genomes. However, in *Ca.* Kuenenia stuttgartiensis the oxidative branch of the TCA cycle is likely mediated by a *Re*-citrate synthase, which operates incompletely to synthesize alpha-ketoglutarate similar to other anaerobic bacteria [11]. The groundwater genomes nzgw511-514 also encoded a *Re*-citrate synthase that was highly similar to the *Re*-citrate synthase identified in *Clostridium kluyveri* (55 - 58%) [12].

**Supplementary Figures**

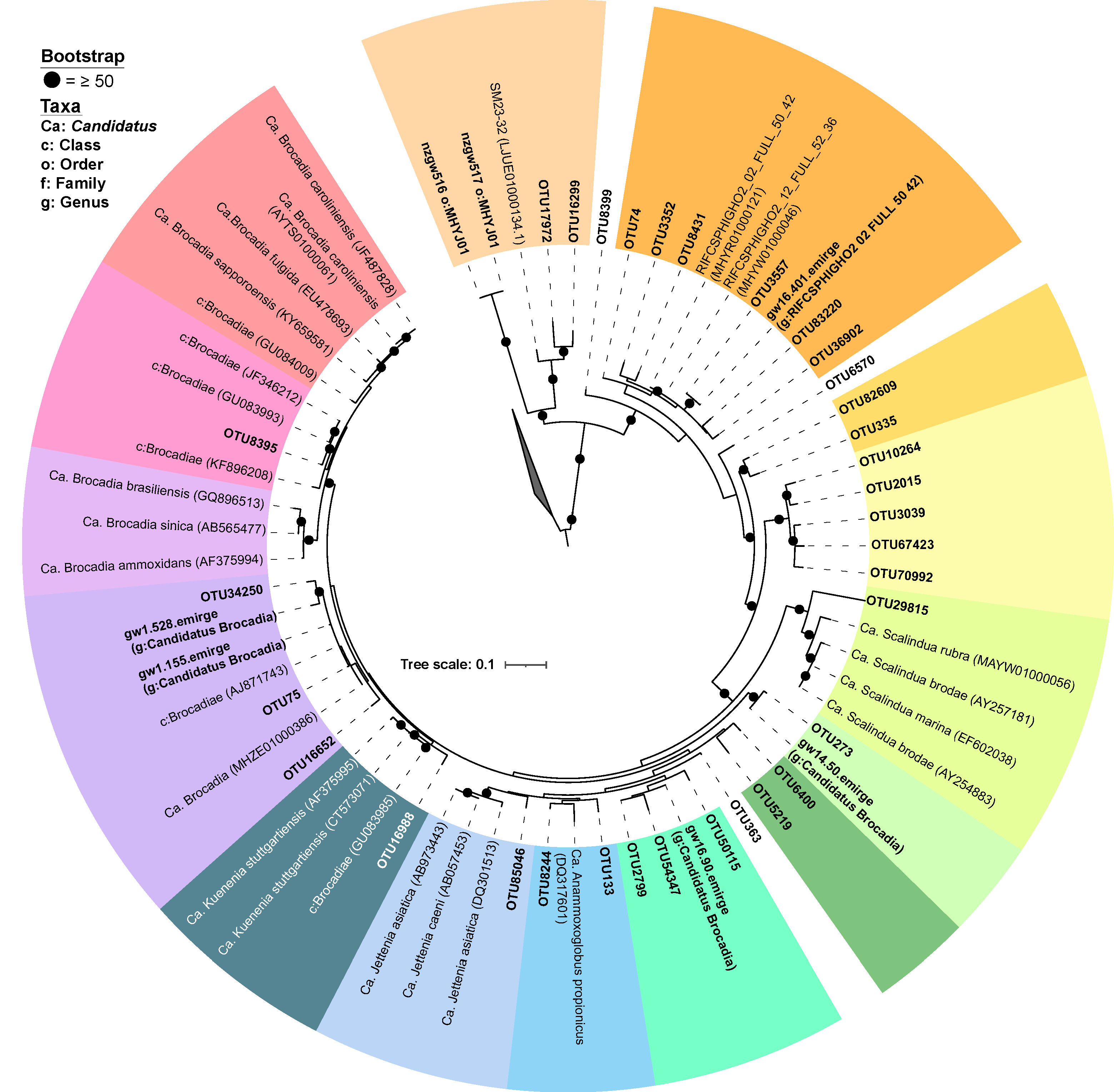

**Figure S1**. Maximum-likelihood phylogenetic tree showing 16S rRNA gene sequences classified as the class *Brocadiae*. Sequences recovered from groundwater in this study (shown in bold font) comprise 16S rRNA gene amplicon OTUs and (near) full length 16S rRNA gene sequences reconstructed from metagenomic data using EMIRGE or SPAdes with identification by Metaxa2. Reference sequences were obtained from the SILVA SSU (r183.1) database. The tree was built using TIM3+F+I+G4 model of substitution using 1,000 bootstrap replicates and annotated in iTOL. Scale bar represents number of substitutions per site. Coloured tiles distinguish different anammox clades.

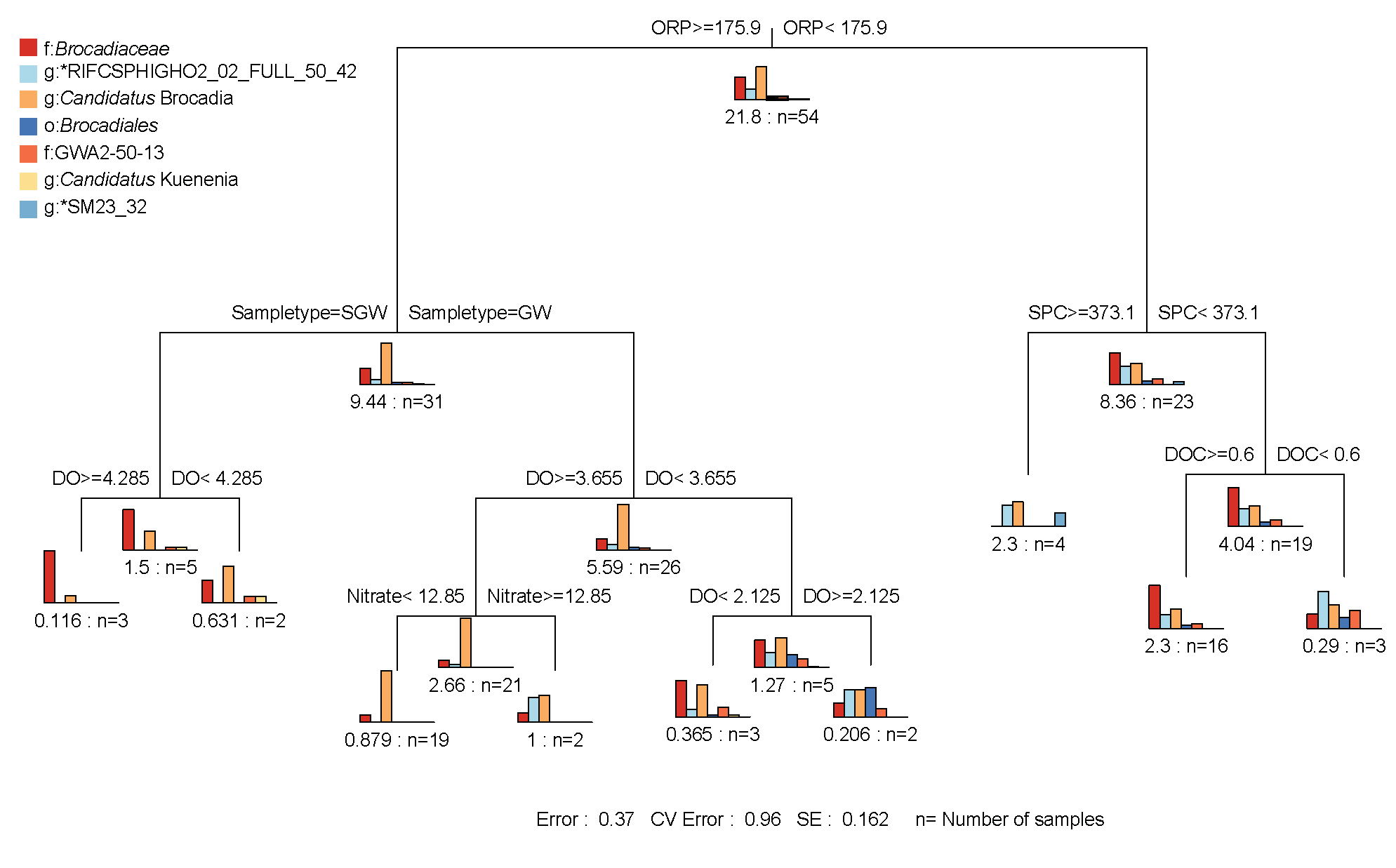

### Figure S2. Multivariate regression tree analysis generated using mvpart v1.6.2 [18] of the relation between abundance of anammox (grouped at lowest level of identification from order *Brocadiales* to genus) and environmental parameters in groundwater samples. ORP values in mV, DO in mg/L, SPC in μScm^-1^, nitrate-N and DOC in g/m^3^. GW = Groundwater, SGW = Sonicated groundwater. * = *Planctomycetes* bacterium. Bar plots show the 16S rRNA gene based relative abundance of anammox bacteria shown in the key.

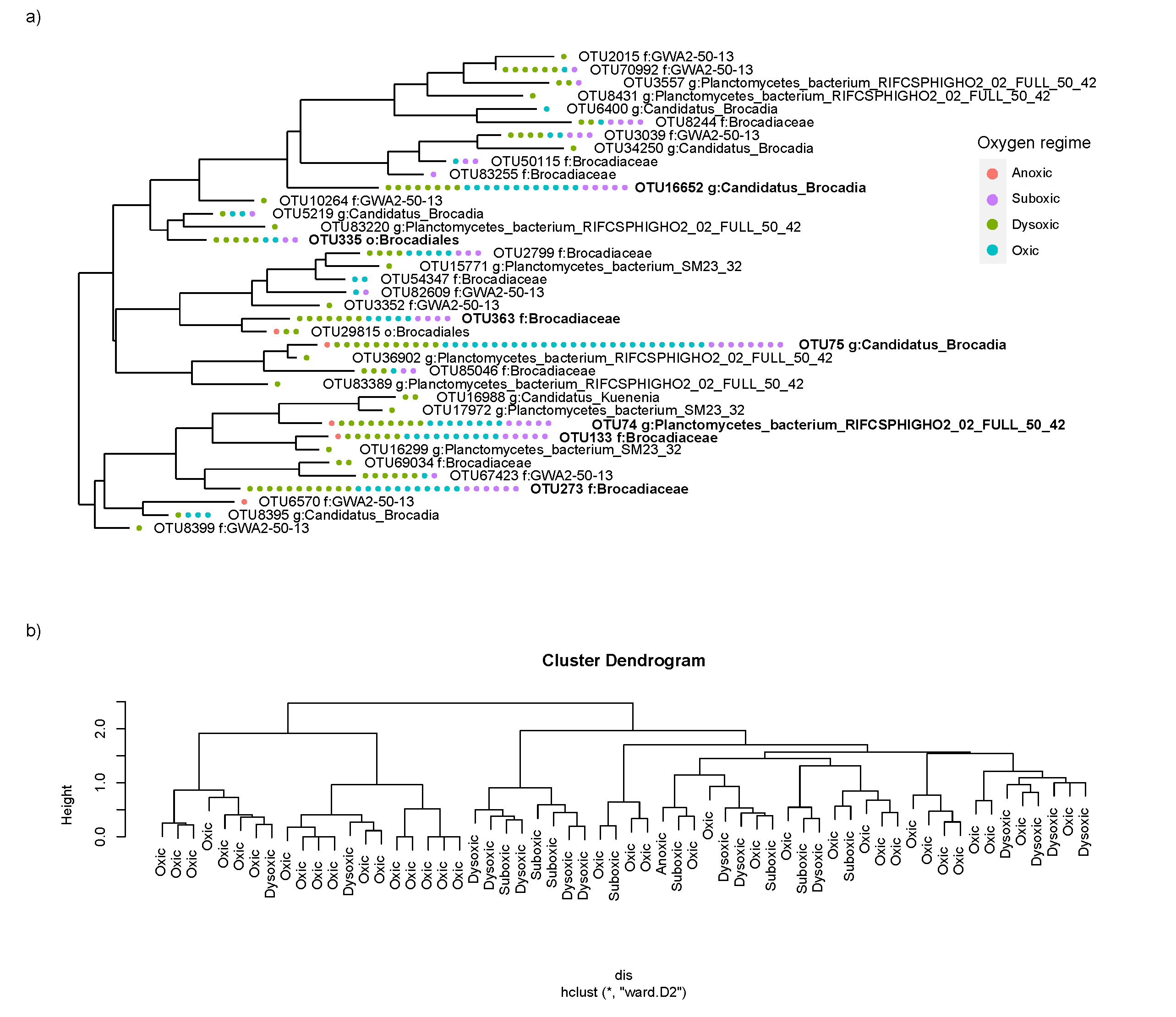

**Figure S3**. **a)** A random fit tree generated using the APE package (RStudio) and plotted with phyloseq using 16S rRNA OTUs of class *Brocadiae* from amplicon data showing presence/absence at sites. Points represent sites and are coloured according to oxygen regime (Anoxic, Suboxic, Dysoxic and Oxic). The 7 most abundant OTUs are shown in bold text. **b)** Cluster dendrogram showing hierarchical cluster analysis on the Bray-Curtis dissimilarity between groundwater samples labelled with oxygen regime and clustered with ward.D2 method.

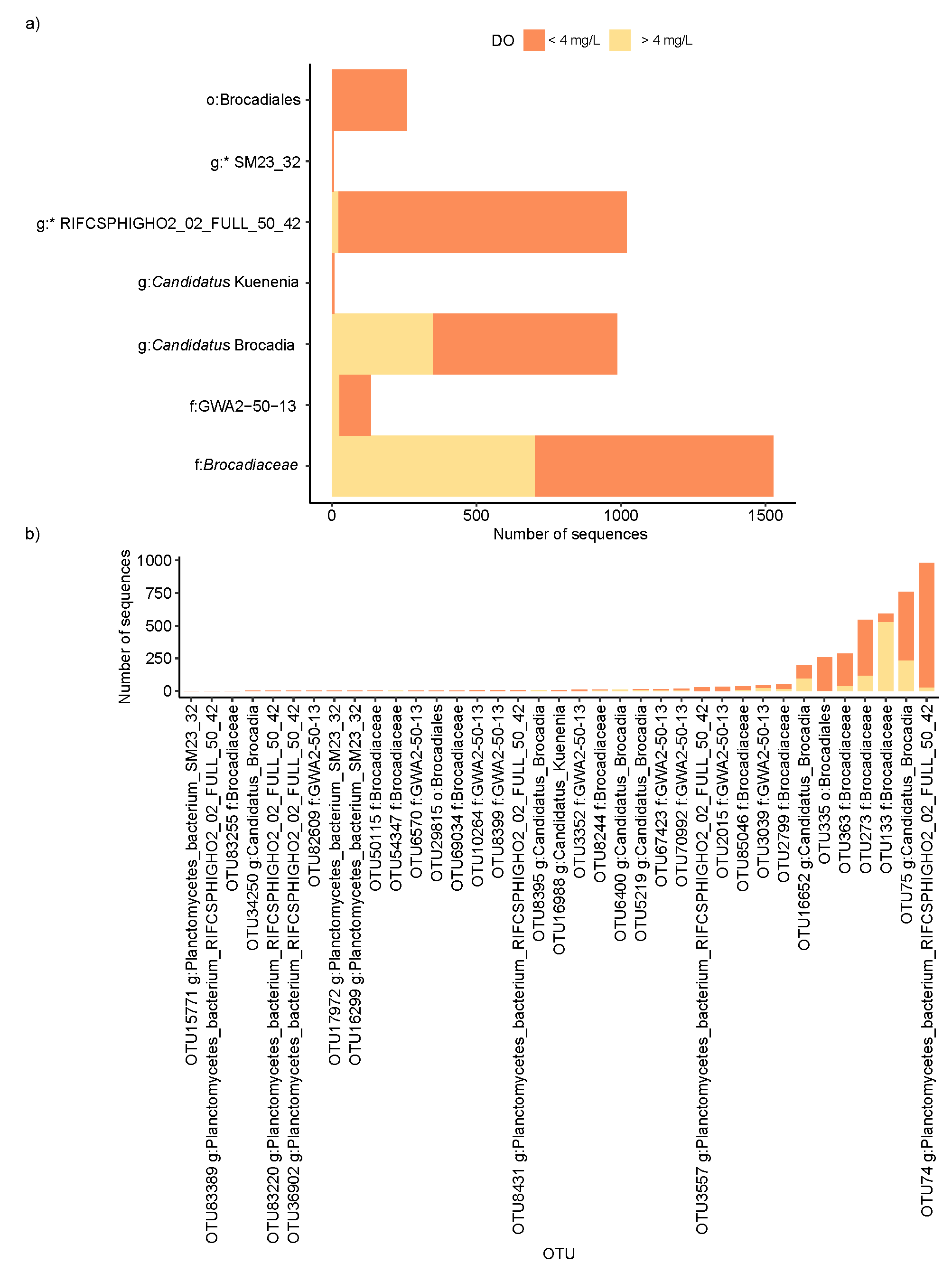

### Figure S4. Phylogenetic composition of the *Brocadiae* communtiy based on 16S rRNA sequences, with oxygen concentration a) Stacked bar plot showing number of sequences from the rarefield OTU table containing anammox (class *Brocadiae*) present in samples > 4 mg/L and < 4 mg/L dissolved oxygen (DO) grouped to the lowest identification level from the order *Brocadiales* to genus level (* = *Planctomycetes* bacterium). b) Stacked bar plot showing the number of sequences from 37 anammox OTUs (class *Brocadiae*) present in samples with > 4 mg/L and < 4 mg/L DO.

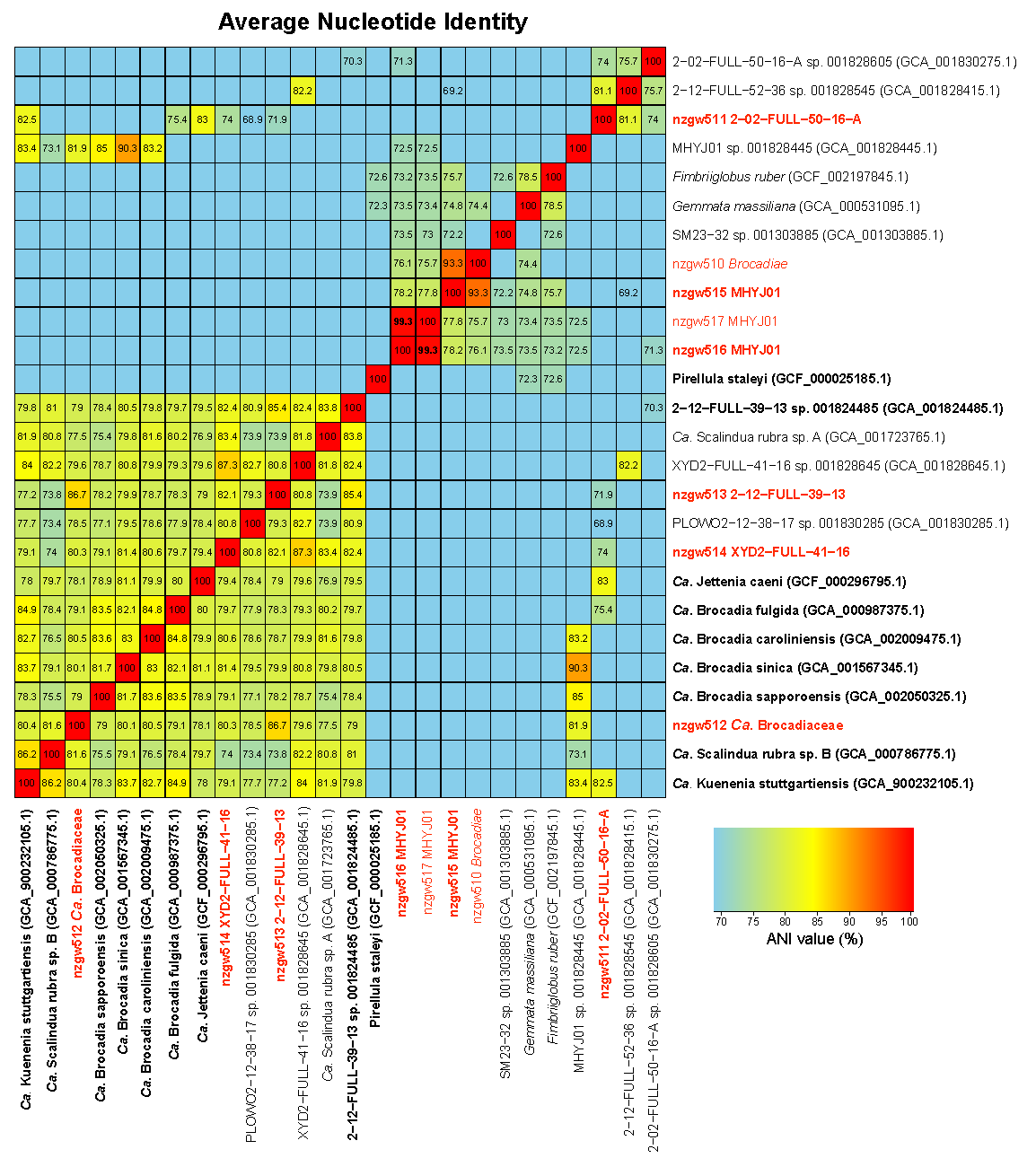

**Figure S5.** Genome similarity heatmap showing pairwise Average Nucleotide Identity (ANI) values of 8 recovered *Brocadiae* genomes from this study (red) and 19 reference genomes (black). Genomes marked with asterisks are non-anammox *Planctomycetes.* Bolded genomes have > 80% estimated completeness and < 5% contamination.

**Figure S6.** Hydrazine synthase subunit B (*hzsB*) abundance (genes and transcripts copies) from 4 wells at sites sampled for metagenomic analysis.

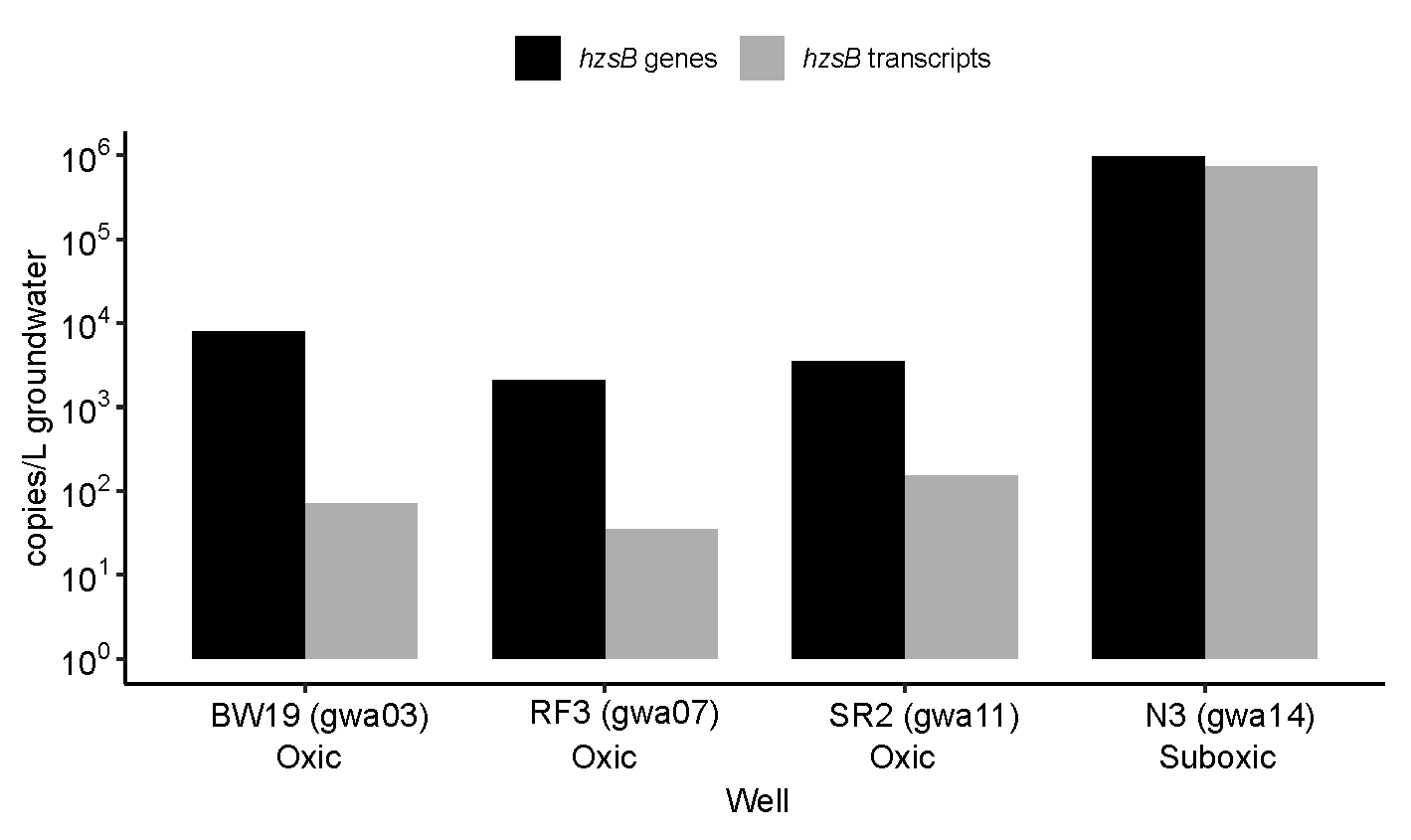
